# Trial-by-trial inter-areal interactions in visual cortex in the presence or absence of visual stimulation

**DOI:** 10.1101/2024.12.05.626981

**Authors:** Dianna Hidalgo, Giorgia Dellaferrera, Giordano Ramos-Traslosheros, Will Xiao, Carlos R. Ponce, Maria Papadopouli, Stelios Smirnakis, Gabriel Kreiman

## Abstract

State-of-the-art computational models of vision focus on fitting trial-averaged spike counts triggered by visual stimuli using overparameterized neural networks. Understanding visual cortex requires predicting the dynamic responses of neurons within and across areas, in single trials, and across different experimental conditions. Here we investigated trial-by-trial inter-areal interactions in the visual cortex by predicting neuronal activity in one area based on activity in another area, including stimulus-driven and non-stimulus-driven activity. We analyzed two datasets: calcium imaging from mouse V1 layers 2/3 and 4, and extracellular neurophysiological recordings from macaque V1 and V4. Neuronal activity could be predicted bidirectionally between L2/3 and L4 in mice, and between V1 and V4 in macaque monkeys, even in the absence of visual stimuli, with distinct contributions for bottom-up and top-down interactions. In mice, we observed a bimodal distribution of neurons, with some neurons primarily driven by visual inputs and others showing predictable activity during spontaneous activity despite lacking consistent visually evoked responses. Predictability depended on the stimulus type, intrinsic neuronal properties, receptive field overlap, and the relative timing of activity across areas. These findings provide initial steps towards computational models that capture the dynamic transformation of neuronal responses throughout visual cortex.

**Impact statement:** Neuronal activity in one visual cortical area can be predicted from another area in single trials, with or without visual stimulation, in both mice and macaques.

## Introduction

To predict the activity of neurons in the visual cortex, multiple studies have focused on correlating external stimuli with trial-averaged responses (***Hubel and Wiesel, 1962; Pasupathy et al., 2020***). Between the stimulus and specific cortical neurons, there is a complex signal processing cascade involving multiple processing stages. Therefore, computational models of visual processing typically gloss over most of the relevant biological machinery in an attempt to fit average firing rates from images (***Serre et al., 2007a; Yamins et al., 2014***).

A mechanistic understanding of the factors that govern firing in the visual cortex requires models that can capture the trial-by-trial transformations across those processing stages. Moreover, neurons throughout the cortex fire “spontaneously” in the absence of any visual input. Thus, by definition, any model predicting neuronal activity that is exclusively dependent on visual stimulation does not account for such fluctuations. Previous studies in mice have revealed significant non-visual influences in neuronal activity in cortex, even in V1, partly accounted for by movement (***Stringer et al., 2019b; Avitan and Stringer, 2022; Polack et al., 2013; Niell and Stryker, 2010; Dadarlat and Stryker, 2017***). These observations contrast with a recent macaque study which did not find the same motor-related spontaneous activity in either V1, V2, or V3 (***Talluri et al., 2023***). Nevertheless, variables that are not related to movement, such as attention, expectation, and arousal, also modulate stimulus- and non-stimulus driven neuronal activity in monkeys (***Reynolds and Chelazzi, 2004; Gazzaley et al., 2007; Okazaki et al., 2008; Gilbert and Li, 2013***), potentially adding to the response variability across stimulus repeats and to neuronal activity in the absence of visual stimuli.

Neuronal interactions between visual areas occur in the presence and absence of visual stimuli (***Chen et al., 2022b; Stringer et al., 2019b; Wosniack et al., 2021; Ringach, 2009; Avitan and Stringer, 2022***). Therefore, such interactions can and should be studied both as a function of sensory inputs and contextual cues but also in the absence of external stimulation or task demands (***Chacron et al., 2003; Hsu et al., 2004; Ringach, 2009***). A paradigmatic example of inter-areal interactions is the series of synaptically-connected laminar (e.g. layer 4 → layer 2/3) and cortical areas (e.g. V1→V2→V4→IT) within the ventral visual stream (***Lee et al., 2016; Felleman and Van Essen, 1991; Markov et al., 2014; Douglas and Martin, 2004; Wang and Burkhalter, 2007; Consortium et al., 2021***). Due to feedforward, feedback, and horizontal connections in the ventral visual stream, the inter-areal interactions could represent reliable shared visual and non-visual information. Several studies examined interactions between visual areas in mice and macaques, focusing on the entire population level (***Semedo et al., 2019, 2022; Tang et al., 2023; Morales-Gregorio et al., 2024***), trial-averaged responses removing transient fluctuations (***Semedo et al., 2019***), neuronal activity in response to only one image presentation (***Semedo et al., 2019, 2022***), or in the absence of any stimulus (***Morales-Gregorio et al., 2024***). Here we investigated interactions between areas in single trials at the level of cortical layers or brain areas across different stimulus types or in the absence of visual stimulation, across different species, and across different recording techniques and temporal resolutions. Throughout, we use the term “interactions” to denote shared variability at the single-trial level, including stimulus-induced correlations and trial-by-trial stimulus-independent fluctuations which can arise from internal state, contextual inputs, or other top-down factors. We focused on multiple simultaneously recorded areas of the ventral visual stream to assess the stimulus- and non-stimulus-driven variability shared between cortical subnetworks. We found that it is possible to reciprocally predict neuronal activity, both during visual stimulation but also during spontaneous activity, and that this predictability depends on the intrinsic properties of each neuron, the degree of receptive field overlap, and the relative timing of activity across areas.

## Results

### Layer 4 activity predicts layer 2/3 activity and V1 activity predicts V4 activity in single trials

We studied neuronal activity from two open datasets: mouse neurons in V1 layer 4 and layers 2/3 (L4 and L2/3; calcium imaging; ***Figure 1***A) (***Stringer et al., 2019a***), and macaque monkey multiunit sites in areas V1 and V4 (extracellular electrophysiology; ***Figure 1***B) (***Chen et al., 2022b***). In addition, we recorded new data from an additional monkey during spontaneous activity and using the same visual stimuli as in ***Chen et al. (2022b***) (Methods). The mouse neuronal recordings we used for this experiment were based on approx. 5,500 neurons per mouse (n=4, ***Table 1***) responding to visual stimuli (drifting gratings or static natural black and white images; total of 7 recording days), in addition to “spontaneous” activity during approximately 30 minutes of gray/black screen presentation on 6 of the 7 recording days. The monkey recordings were based on a range of 32-803 electrodes per animal (3 monkeys, L, A, and D; ***Table 2***) responding to visual stimuli (full-size static checkerboard image; n=3 recordings for monkey L, n=1 for monkey A, n=2 for monkey D), small and thin bar slow-moving in a small clockwise square direction (n=1 for monkeys L and A); or large and thick bar fast-moving in a big clockwise square direction (n=1 for monkeys L and A) in addition to spontaneous activity during gray screen presentation in all recording days (n=5 for monkey L, n=3 for monkey A, n=2 for monkey D). The recordings consisted of multi-unit activity (MUAe) and local field potentials (LFP) in monkeys L and A retrieved from the open dataset by Chen et al. (***Chen et al., 2022b***). Monkey D recordings were acquired using the same checkerboard stimulus presentation conditions as stated in ***Chen et al. (2022b***). There was also a lights-off condition, where the head-fixed monkeys were free to open or close eyes for approximately 10-25 minutes (monkeys L/D; 2-3 recording days). We omitted channels with signal-to-noise ratio of less than 2, or channels that were considered “spurious” by the authors of the open dataset (***Chen et al., 2022b***).

**Table 1.**
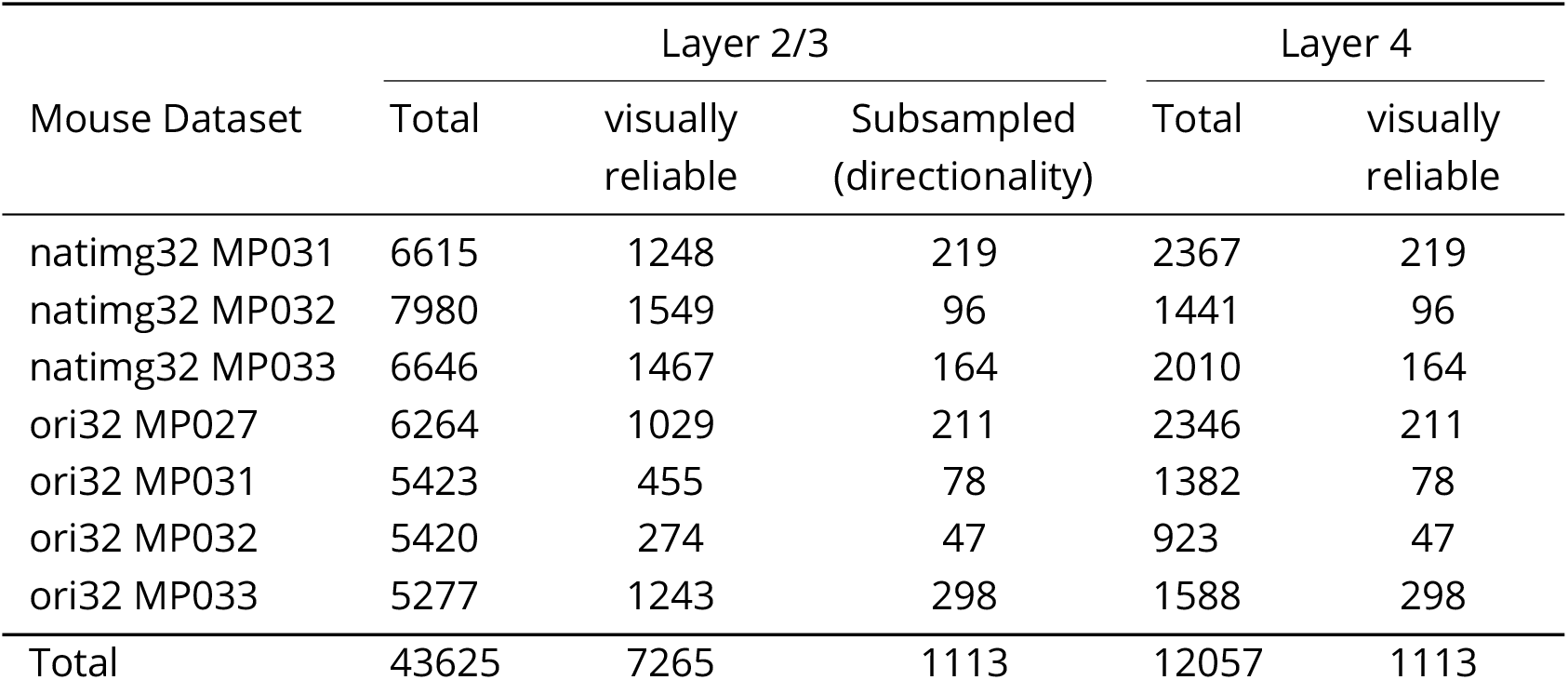
Mouse neuron counts used for inter-layer prediction and analyses. A total of 7 recordings were used to perform prediction experiments. Each row corresponds to a recording day, containing the dataset recording type (Mouse Dataset), total number of neurons, and visually reliable neurons (see Methods). Fourth column: In the directionality prediction experiments, the area containing more neurons (L2/3) was further subsampled to match the number of L4 neurons. The dataset recording type names contain either “ori32” or “natimg32”, in addition to the mouse name (MP0-). “natimg32” represents the dataset of the 32 natural image presentations. “ori32” represents the dataset of the 32 drifting gratings.

**Table 2.** Monkey site counts used for inter-cortical prediction and analyses. First column: First letter denotes the monkey name, followed by the date, followed by the dataset type acquired during the session (C: Checkerboard presentations, G: Gray screen, L: Lights-off condition, LB: Large, thick moving bars, SB: Small, thin moving bars). Fourth column: In the directionality prediction experiments, the area containing more sites was further subsampled to match the number of V4 sites.

| Monkey Datasets | V1 |  |  | V4 |  |
| --- | --- | --- | --- | --- | --- |
|  | Total | visually reliable | Subsampled (directionality) | Total | visually reliable |
| L 090817 (C,G,L) | 627 | 555 | 73 | 96 | 73 |
| L 100817 (C,G,L) | 688 | 587 | 83 | 115 | 83 |
| L 250717 (C,G,L) | 645 | 537 | 72 | 86 | 72 |
| L 260617 (LB,G) | 645 | 592 | 82 | 86 | 82 |
| L 280617 (SB,G) | 645 | 518 | 68 | 86 | 68 |
| A 041018 (C,G) | 571 | 357 | 44 | 76 | 44 |
| A 280818 (LB,G) | 571 | 381 | 58 | 76 | 58 |
| A 290818 (SB,G) | 571 | 251 | 30 | 76 | 30 |
| D 250225 (C,G,L) | 16 | 8 | - | 16 | 7 |
| D 260225 (C,G,L) | 16 | 10 | - | 16 | 10 |
| Total | 4995 | 3796 | 510 | 729 | 527 |

**Figure 1.**
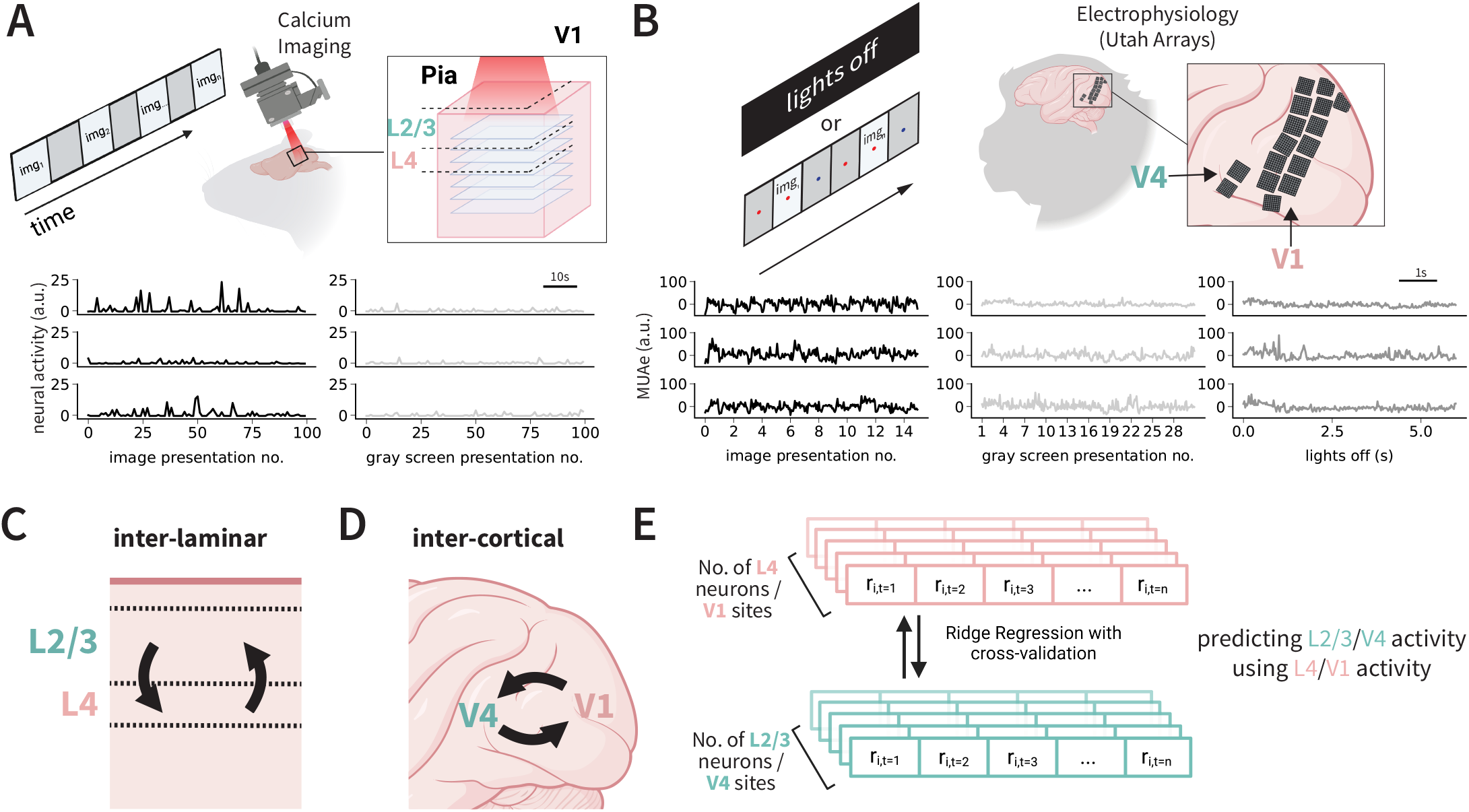
Predicting trial-to-trial and timepoint-to-timepoint neuronal activity between areas. **A**. Top: Experimental set-up to record two-photon Calcium imaging activity data from layers 2/3 (L2/3) and layer 4 (L4) in rodent V1 upon presentation of gratings, natural stimuli or gray screen images (represented as *img*_*n*_) (***Stringer et al., 2019a***). Deconvolved calcium imaging traces were z-scored using baseline activity during 30 minutes of gray screen presentation before/after image presentation (***Table 1***). Bottom: Sample z-scored neuronal activity from 3 different neurons in response to 100 presentations of drifting gratings (left) or gray screen presentations (right). Each activity value corresponds to one image presentation, and was calculated as the average of two calcium imaging video frames (666 ms or 800 ms, see details in Methods). B.Top: Experimental set-up for the neuronal activity data from monkeys V1 and V4 (***Chen et al., 2022b***). Electrophysiological activity was simultaneously recorded across 1,024 channels from 16 Utah arrays (***Table 2***). Bottom: Envelope multiunit spiking activity (MUAe) from 3 different sites in response to multiple presentations of a repeated 400 ms full-field checkerboard image (left, baseline mean-subtracted), 200 ms gray screen (middle), or during a lights-off condition (30 minutes total; right). Each value corresponds to aggregated MUAe activity in a 25-ms bin. B. Overview of inter-laminar relationships examined in mouse V1. “Lower level” layer 4 (L4) neuronal activity is used to predict “higher level” layer 2/3 (L2/3) activity and vice versa. **D**. Overview of inter-cortical relationships examined in monkeys, where lower-level V1 is used to predict higher-level V4 (blue arrow) and vice versa (red arrow). **E**. Illustration of linear ridge regression method used for inter-areal prediction. Neuronal activity in response to presentation number *i* (labeled “*r*_*i*_”) at time *t* from one area (e.g., mouse V1 L2/3 or monkey V1) was used to predict activity in the other area (e.g., mouse V1 L4 or monkey V4) (***Semedo et al., 2019***). Predictability was evaluated using 10-fold cross-validation across presentation trials in mice, and across 25-ms timepoints in monkeys (Methods).

We evaluated two types of interactions between areas: inter-laminar (***Figure 1***C; mouse V1) and inter-cortical (***Figure 1***D; monkey). The intention of using data from mice and monkeys was *not* to compare the two species but rather to study the interactions between areas *within* each species. We used linear ridge regression to predict neuronal activity in one area from activity in the other area in single trials (***Figure 1***E, see Methods) (***Semedo et al., 2019***). For each neuron/site in the target area we *fit* a linear readout from the other area’s simultaneous population activity on the same trial, with an *L*_2_ (“ridge”) penalty. Performance was evaluated using cross-validation over trials and quantified as squared Pearson’s r (hereafter, “explained variance” or EV, Methods). ***Figure 2***A shows sample neuronal activity from three example mouse V1 L2/3 cells during image presentation (black traces). Overlaid, the figure also shows the predicted neuronal activity (red). The predicted neuronal activity is shown as a function of the actual activity in response to every image presentation for the same example cells in ***Figure 2***C. The top cell illustrates a case where the predicted activity closely matches the actual activity (*EV* = 0.67), the middle cell shows a typical case (*EV* = 0.39), and the bottom cell illustrates a case where the predictions deviated from the actual neuronal activity (*EV* = 0.07). We focused on neurons deemed “visually reliable” (~17% of total L2/3 neurons; ***Table 1***, Methods, see results for all neurons in ***Figure 2—Figure Supplement 1***C). The ridge regression model predicted single-trial L2/3 activity from L4 activity across both types of visual stimuli with an average EV of 0.28 ± 0.16 (mean ± stdev. across neurons, ***Figure 2***E) whereas the shuffle control mean EV was 0.004 ± 0.002.

**Figure 2.**
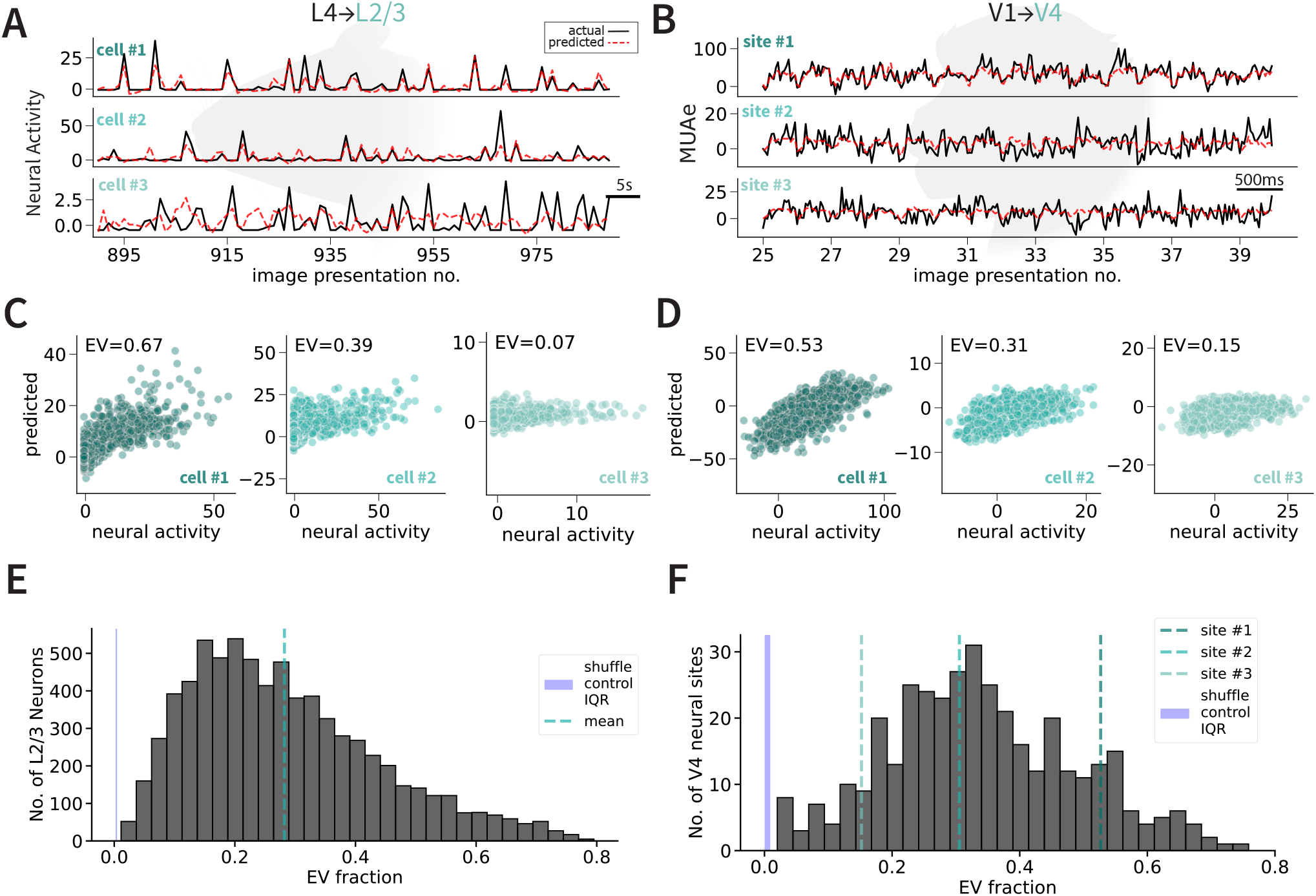
Lower level activity can predict higher level activity in both rodent and primate brains. **A**. Example neuronal activity (z-scored, black) in response to stimulus presentations (drifting gratings) in mouse V1 L2/3 along with regression-model predictions (red) for a typical cell (2, middle), cell in the top 10% percentile of predictability (1, top), and bottom 10% percentile (3, bottom). **B**. Same as A for monkey MUAe activity in response to a full-field checkerboard image in three V4 neuronal sites. **C**. Predicted neuronal activity versus actual neuronal activity in response to stimuli for the mouse L2/3 cells 1, 2, and 3 shown in A. Each point represents 800 ms, corresponding to a stimulus presentation. *r* values (top left) indicate the correlation coefficient. **D**. Same as C for monkey V4 neuronal sites 1, 2, and 3. Each point represents one 25-ms timepoint during the 400-ms presentation. **E**. Distribution of EV fraction in L4→L2/3 regressions of neural activity in response to visual stimuli in cells that were deemed visually reliable in 4 mice and 7 recording days (*n* = 7, 265 neurons, Methods). Performance using 10-fold cross-validation across trials was quantified as squared Pearson’s r, referred to as explained variance (EV) fraction. The three vertical lines show the 3 examples in part **A, C**. The blue solid shaded rectangle (here and throughout) represents the interquartile range (IQR) shuffle control performance, where the activity timepoints of one area were randomly shuffled. **F**. Distribution of EV fraction in V1→V4 regressions of neural activity in response to visual stimuli in sites deemed visually reliable in monkey L (5 recording days, 68–82 V4 sites recorded per day; *n* = 378 total site recordings).

In monkeys, trial-to-trial variations in V4 activity were predicted from V1 activity across the three types of visual stimuli. Example recording sites for monkey L are shown in ***Figure 2***B, D (see also ***Figure 2—Figure Supplement 1*** D–G for examples from two additional monkeys, A and D). The ridge regression model predicted the single-trial responses in V4 activity from V1 activity with an average EV of 0.30 ± 0.15 (***Figure 2***F for monkey L, see ***Figure 2—Figure Supplement 1***H for monkey A and ***Figure 2—Figure Supplement 1***I for monkey D), whereas the shuffle control mean EV was 0.005 ± 0.005. Most sites in V4 (72.3%) were deemed visually reliable; EV results for all monkey sites are shown in ***Figure 2—Figure Supplement 1***J). EV performance varied across the three monkeys (*EV*_*L*_ = 0.34, *EV*_*A*_ = 0.21, *EV*_*D*_ = 0.11). This variation may be due to predictor population size (***Table 2***). We subsampled sites in monkey L to match the numbers of monkey A and monkey D (20 permutations; ***Figure 2—Figure Supplement 1***L; subsampling was performed for all analyses hereafter unless noted). Subsampled monkey L matching the number of sites in monkey A (monkey *L*_*A*_) showed an average EV of 0.35 ± 0.14, statistically higher compared to the EV for monkey A (*p* < 0.001, permutation test). Similarly, subsampled monkey L matching the number of sites in monkey D (monkey *L*_*D*_) showed an average EV of 0.19 ± 0.11, statistically higher compared to the EV for monkey D (*p* < 0.001, permutation test).

We asked whether the conclusions were dependent on the model and parameter choices. As an alternative approach to ridge regression, we also fit a Poisson general linear model (GLM) enforcing non-negativity to monkey MUAe without baseline subtraction, using identical folds, bin widths, and temporal gaps as in the ridge regression analysis. The results from the Poisson GLM were consistent with the results obtained using ridge regression (***Figure 2—Figure Supplement 1***M). Additionally, to evaluate the impact of the bin size on the results, we re-estimated EV after binning MUAe into 10–200 ms. EV increased with bin size and remained well above shuffle controls across conditions (***Figure 2—Figure Supplement 2***A–C, *p* < 0.001, paired permutation test), indicating that inter-areal predictability is robust across a broad range of physiologically relevant timescales.

In sum, it was possible to provide estimates of neuronal activity in single trials in both species, across different layers within primary visual cortex in mice and across different visual cortical areas in monkeys.

### Inter-cortical predictions are asymmetrical

In the previous section, we demonstrated the possibility of predicting L2/3 activity from L4 activity and V4 from V1. We asked whether we could also predict neuronal responses in the opposite direction. We could predict L4 activity from L2/3 activity in mice and V1 activity from V4 activity in monkeys (***Figure 2—Figure Supplement 1***A for mice, ***Figure 2—Figure Supplement 1***K for monkeys). To directly compare predictability between directions *within* each species, we matched the number of predictors (i.e., number of neurons/sites used to predict activity) and the degree of self-consistency (split-half *r* values) by randomly subsampling in each layer or cortical region prior to computing the predictability metrics (***Figure 3***A, C, Methods).

**Figure 3.**
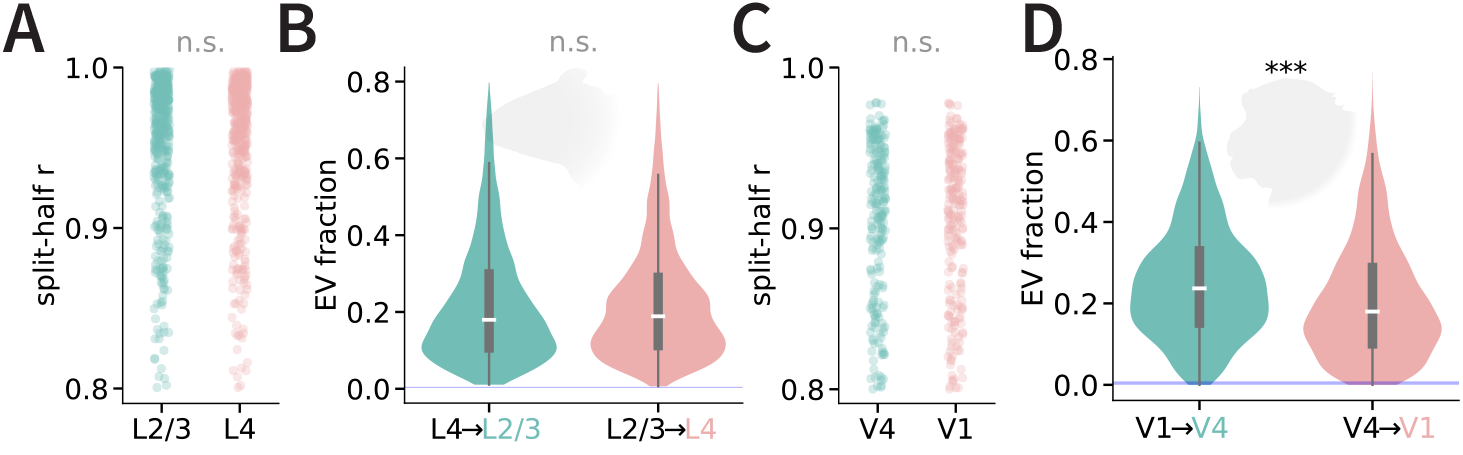
Asymmetry in inter-cortical predictability in monkeys but not inter-laminar predictability in mice. **A**. Split-half reliability (Methods) for the *n* = 298 neurons (per area) in mouse MP033 drifting gratings presentation recording of V1 L2/3 (green) and L4 (coral) used to perform directionality comparisons. Neurons were randomly subsampled to match the numbers and self-reliability in the two distributions. Here and throughout, asterisks indicate statistically significant differences using a hierarchical independent permutation test (10,000 permutations): * *p* < 0.05, ** *p* < 0.01, *** *p* < 0.001; “n.s.” indicates *p* > 0.05. **B**. Violin plots describing the distribution of EV fraction for L4→L2/3 (green) and L2/3→L4 (coral) predictions across all 7 stimulus recordings (*n* = 1, 113 neurons per area). Violin plots (here and throughout) represent the distribution of neuron/site values, with width representing density and inner boxplot representing the interquartile range. Whiskers of each inner box represent the data range. **C**. Example of split-half reliability for the *n* = 74 sites (per area) in monkey L checkerboard recording (date=090817) of V4 (green) and V1 (coral) used to perform directionality comparisons. **D**. Violin plots describing the distribution of EV fraction for V1→V4 (green) and V4→V1 (coral) across all 5 stimulus recordings (*n* = 786 sites recordings per area).

In mice, there was no statistically significant difference between L4→L2/3 and L2/3→L4 directions (*p* = 0.69, hierarchical permutation test, ***Figure 3***B). In monkeys, after matching predictor count and split-half correlation values, the EV fraction in the V1→V4 direction was higher than in the V4→V1 direction (*p* < 0.001 for monkey L, ***Figure 3***D; *p* = 0.01 for monkey A, ***Figure 3—Figure Supplement 2***C). This directional asymmetry was stable across temporal bin sizes in monkey L (***Figure 2—Figure Supplement 2***D). In monkey A, the asymmetry was small and was not robust across different bin sizes (***Figure 2—Figure Supplement 2***E), consistent with the noisier data and weaker effect size in that animal. There were not enough sites with similar reliability distributions in monkey D to perform the matched subsampling directionality tests.

Although we accounted for predictor size and reliability distributions, differences in prediction performance between directions could still reflect additional inherent differences in target-population properties (***Figure 3—Figure Supplement 1***F–J). To control for these differences, we modeled EV as a function of target–population properties and used the residuals for inference, 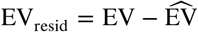. Covariates included self-consistency, SNR, one–vs–rest *r*^2^, and variance metrics (Methods). After adjustment, the difference between inter-cortical directions remained statistically significant overall (*p* < 0.001 for monkey L, *p* < 0.001 for monkey A, ***Figure 3—Figure Supplement 2***F– G). These results suggest that the differences between directions cannot be attributed to target-related neuronal properties. We also analyzed the residuals in mice prediction directions, and after controlling for neuron properties, we still found no statistically significant difference in predictability directions (***Figure 3—Figure Supplement 2***E, *p* = 0.22 for stimulus activity, *p* = 0.48 for gray screen activity).

### Predictability of neuronal activity is dependent on the visual stimulus

We evaluated whether the predictability of neuronal activity varied with the type of visual stimulus presented to the animal. This analysis was done *within* each species, and should not be interpreted as comparison between mice and macaques. In mice, we compared the inter-laminar prediction of neuronal activity of visually reliable neurons in response to drifting gratings versus natural images (***Figure 4***A). We could predict mouse L4 and L2/3 activity under both stimulus conditions (*p* < 0.001, paired permutation test of prediction vs. shuffled frames prediction). Predictability was higher for drifting gratings than natural images in the L4→L2/3 direction (***Figure 4***B; *p* < 0.001, hierarchical permutation test).

**Figure 4.**
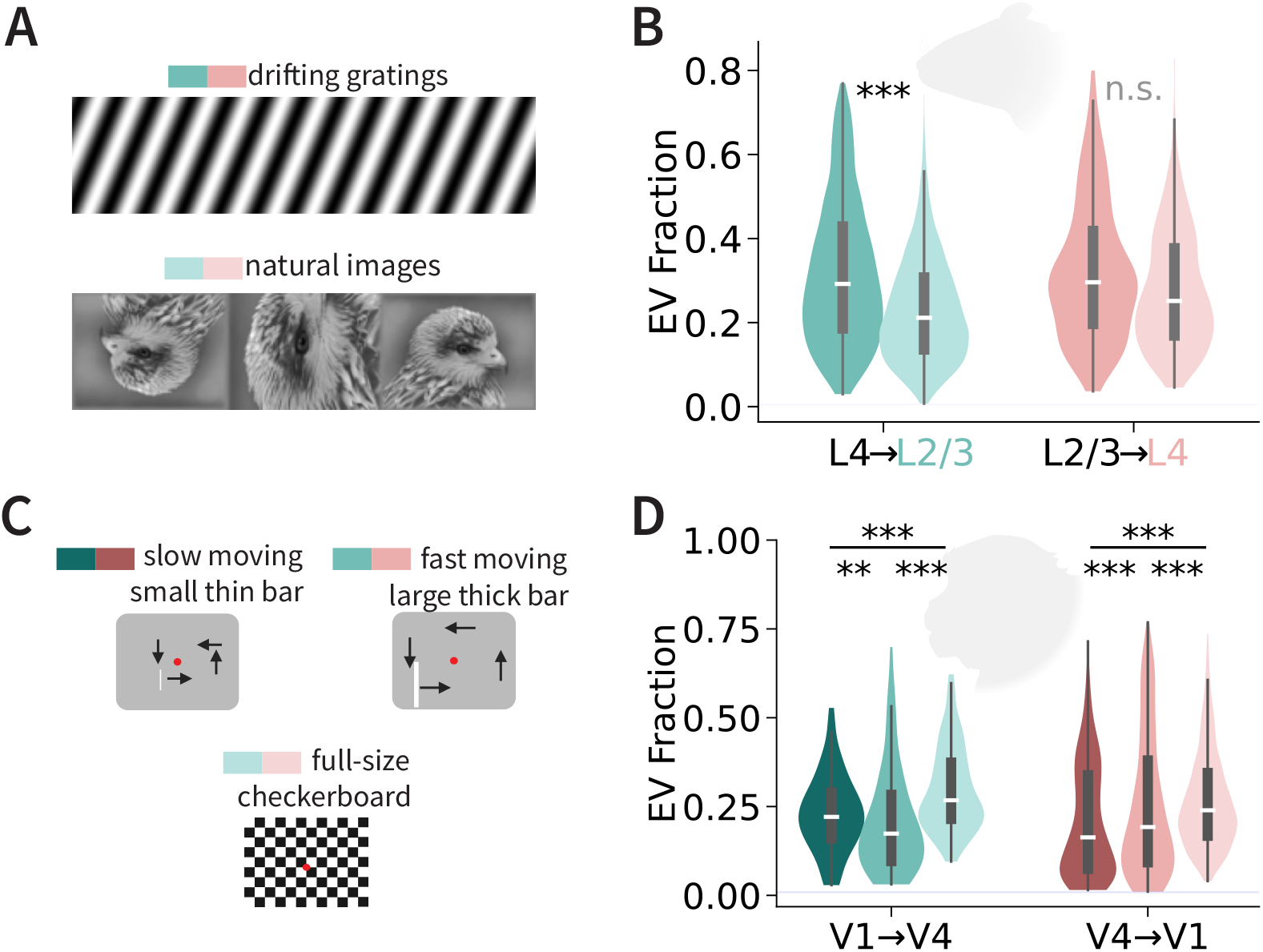
Stimulus type influences neuronal predictability. **A**. Illustration of the two types of stimuli (drifting gratings and static natural images) presented to the mouse during calcium imaging. **B**. Across-layer predictability in mouse V1 for each stimulus type (dark: drifting gratings, light: natural images) and prediction direction. **C**. Illustration of the three types of stimuli presented to the monkeys (***Chen et al., 2022b***). The slow-moving, small, thin bar moved near the fixation point for 1 s in each of the four directions, while the fast-moving, large, thick bar moved towards the edges of the screen for 1 s in each of the four directions. The full-field checkerboard image was presented repeatedly (400 ms each presentation). **D**. Across-area predictability for each stimulus type (dark: slow bars, medium: fast bars, light: checkerboard) and direction.

In monkeys L and A, we compared inter-cortical predictability of visually reliable sites in responses to a slow-moving small thin bar, fast-moving large thick bar, and a full-size checkerboard image (***Figure 4***C). As expected, V1 and V4 could predict each other’s neural activity across all stimulus types (*p* < 0.001, paired permutation test of prediction vs. shuffled frames prediction). The predictability was highest in both directions for neuronal activity in response to a full-field checker-board image (monkey L ***Figure 4***D; monkey A ***Figure 4—Figure Supplement 1***F). In the V1→V4 direction, the EV fraction was higher when predicting a slow-moving small thin bar compared to a fast-moving large thick bar (monkey L ***Figure 4***D, left; monkey A ***Figure 4—Figure Supplement 1***F, left).

In the V4→V1 direction, there was no consistency in prediction differences between monkey L and monkey A. In monkey L (V4→V1), the EV for fast-moving large thick bars was larger than for the slow-moving thin bars (*p* < 0.001, permutation test; ***Figure 4***D, right), whereas this difference was flipped in monkey A (*p* < 0.001; ***Figure 4—Figure Supplement 1***F, right). This observation could be due to differences in the degree of reliability in the neuronal responses between monkeys (***Figure 4—Figure Supplement 1***H, L for monkeys L, A, respectively). We modeled EV as a function of SNR, self-consistency (split-half *r*), variance across time within stimulus, and variance across trials within timepoint, and used 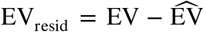 for inference (Methods). In monkey L (V4→V1), the difference in EV residuals between thin and thick bars was not statistically significant (*p* = 0.44; ***Figure 4—Figure Supplement 1***K, right), whereas this difference remained statistically in monkey A (*p* < 0.001; ***Figure 4—Figure Supplement 1***O, right). Lastly, in the V1→V4 direction, EV residuals remained statistically higher when predicting the responses to the thin bar compared to the thick bar in both monkeys L and A (*p* < 0.001 for both monkeys; ***Figure 4—Figure Supplement 1***K,O, left).

### Neuronal activity could be predicted even during spontaneous activity

Given the dependence on the visual stimulus, we next asked whether it would be possible to predict neuronal responses in the absence of any visual stimulus, during “spontaneous activity”. We studied this dependence separately *within* each species. We compared the predictability of stimulus-evoked activity in mice (drifting gratings and natural images) versus the predictability of activity recorded during gray screen presentations. This comparison was conducted in both visually (SNR >2 & split-half *r* >0.8) and non-visually (SNR <2 & split-half *r* <0.8) reliable neurons (n=3 mice; mouse MP027 did not undergo 30 min. of gray screen presentation). In visually reliable neurons, neuronal activity could still be predicted during the gray screen condition (*p* < 0.001, hierarchical paired permutation test compared with shuffled frames). There was a significant reduction in EV during gray screen compared to visual stimulus presentation (***Figure 5***A left, *p* < 0.001, hierarchical paired permutation test). In non-visually reliable neurons, predictability was higher during the gray screen condition compared to stimulus presentation (***Figure 5***A right, *p* < 0.001, hierarchical paired permutation test).

**Figure 5.**
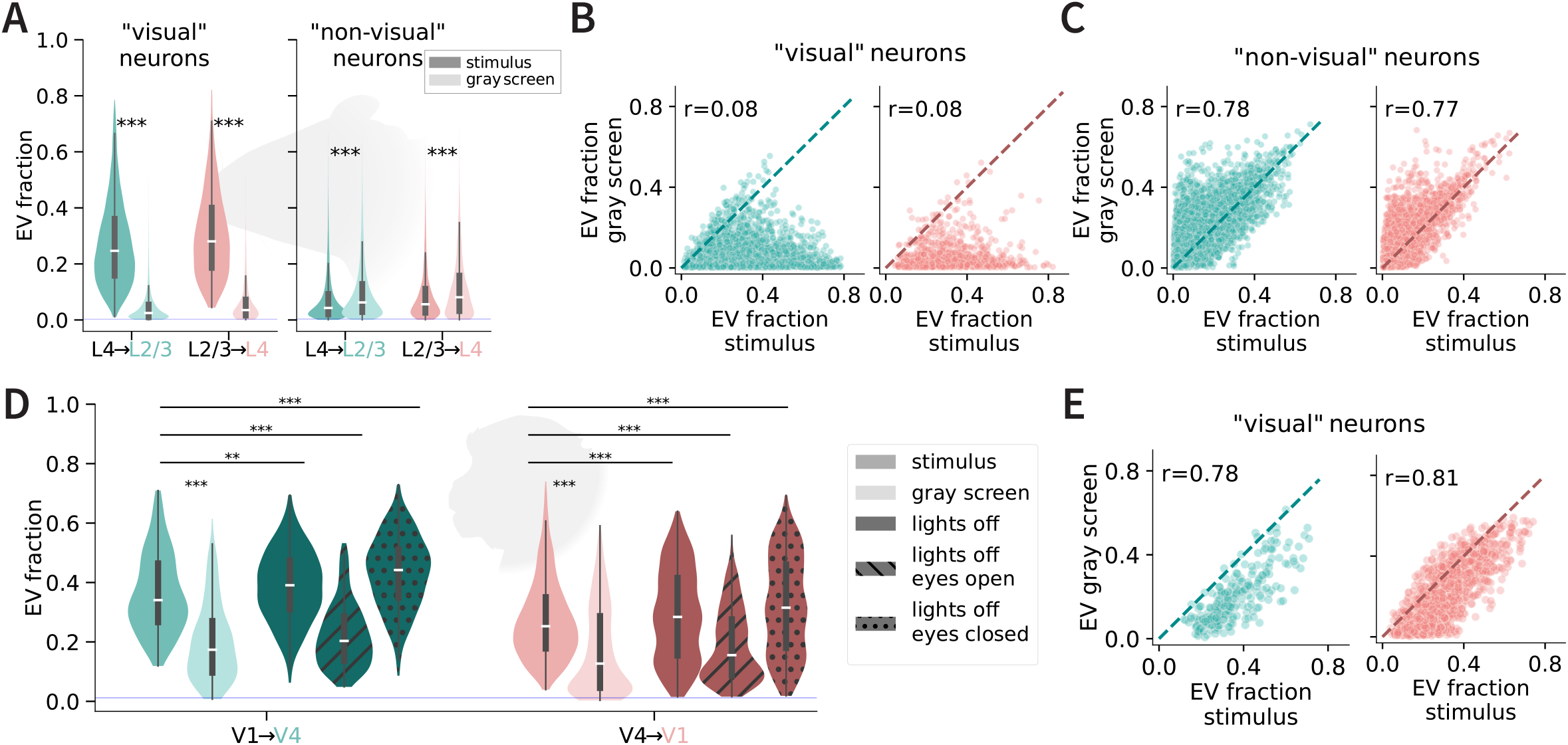
Spontaneous activity can also be predicted. **A**. EV fraction of neuronal activity in response to stimulus presentation (dark violins) or gray screen presentation (light violins) for neurons deemed visually (left) or non-visually (right) reliable (Methods). **B**. Correlation between EV in responses to gray screen (y-axis) versus stimulus presentation (x-axis) in mouse V1 visually reliable neurons (L4→L2/3:left, green; L2/3→L4: right, coral). The diagonal line represents the line of equality (y=x). *r* is the Pearson’s r coefficient. **C**. Same as **B**, but for non-visually reliable neurons. B. EV during stimulus presentations (checkerboard image, green), gray screen presentations (light green), or during lights off (dark green). The lights-off condition is further separated into periods when the eyes were open or closed. All lights-off conditions were sub-sampled (10 permutations) to contain similar training lengths as the stimulus and gray screen presentation recordings. **E**. Correlation between EV in responses to gray screen (y-axis) versus stimulus presentation (x-axis) in monkey visually reliable neurons (V1→V4:left, green; V4→V1: right, coral). The diagonal line represents the line of equality (y=x). *r* is the Pearson’s r coefficient. All recorded sites were pulled from the 3 recording days of checkerboard presentations.

While there was no correlation between neuronal predictability in the responses to visual stimulus presentations and in the response to gray screen presentations in visually reliable neurons (***Figure 5***B), there was a strong correlation for non-visually reliable neurons (***Figure 5***C). The difference in predictability in the absence of a stimulus (gray screen presentations) could in principle change according to the directionality in inter-laminar interactions. However, there was no statistically significant difference in the EV fraction between laminar directions (L4→L2/3 vs. L2/3→L4), in the same “visually reliable” group and even when subsampling “non-visual” neurons (***Figure 3***— ***Figure Supplement 2***A).

In monkeys, we only focused on visually reliable sites since the majority of the neuronal population was visually reliable (***Figure 2—Figure Supplement 1***J). Additionally, an SNR of less than 2 (one of the requirements to define non-visual neurons in the mouse data) most likely reflects artifactual issues with the electrode recording the multiunit site (***Chen et al., 2022b***). We compared inter-areal prediction of stimulus presentation activity (all monkeys for checkerboard presentation; monkeys L and A for moving bars), gray screen presentation (all monkeys), and during lights-off (monkeys L and D). The predictability of neuronal activity in response to gray screen presentation remained statistically above chance (*p* < 0.001 paired permutation test of prediction vs. shuffled frames prediction for all monkeys). The predictability of neuronal activity in the absence of visual stimulus was lower compared to checkerboard presentations (***Figure 5***D, *p* < 0.001 for monkey L; ***Figure 5***— ***Figure Supplement 1***E, *p* < 0.001 for monkey A; ***Figure 5—Figure Supplement 1***F left, *p* < 0.01 for monkey D). The same conclusions held when comparing moving bar vs. gray screen presentations (***Figure 5—Figure Supplement 1***D for monkey L; ***Figure 5—Figure Supplement 1***E for monkey A). Intriguingly, the EV fraction in the lights-off condition was statistically higher than during the stimulus presentations in both directions in monkey L, whereas the EV fraction in the lights-off condition in monkey D was lower (***Figure 5—Figure Supplement 1***F). Eye closure and sleep can induce global oscillations (***Hohaia et al., 2022***) and therefore may lead to correlated neuronal activity, causing an increase in predictability. Monkey L spent a significant portion with its eyes closed (***Chen et al., 2022b***), unlike monkey D, who spent almost the entire time with its eyes open (eyes closed duration < 3*s* in both recording days). To test the effects of eye closure, we further separated the lights-off neuronal activity of into periods where the eyes of monkey L were open or closed. The EV fraction was statistically significantly higher than EV fraction during stimulus presentation activity only during the eyes-closed period (***Figure 5***D).

The correlation in visual predictability between stimulus presentation and spontaneous activity was high across all types of spontaneous conditions (***Figure 5***E and ***Figure 5—Figure Supplement 1***C for monkey L; ***Figure 5—Figure Supplement 1***I, J for monkeys A, D, respectively). When assessing the inter-cortical prediction directionality during spontaneous conditions, in monkey L, we found the same asymmetrical relationship as in ***Figure 3***, where V1→V4 EV fraction was significantly higher than V4→V1 prediction in both gray screen (*p* < 0.001, permutation test) and lights-off (*p* < 0.001, permutation test) conditions (***Figure 3—Figure Supplement 2***B). In monkey A gray screen presentation activity, we did not find any statistically significant difference between direction predictions (***Figure 3—Figure Supplement 2***C, right). However, when accounting for neuronal property differences between the areas, we found the same direction EV asymmetry across both monkeys (*p* < 0.001; ***Figure 3—Figure Supplement 2***F for monkey L; ***Figure 3—Figure Supplement 2***G for monkey A).

### Behavioral metrics account for only a small portion of predictability

Because inter-areal predictability persists in the absence of visual drive, we asked whether behavioral signals could partly account for the inter-laminar and inter-cortical interactions. In mice, augmenting the neural predictability model with face–motion singular value decomposition (SVD) components and running speed did not significantly increase EV relative to the neural-only model in the L2/3 to L4 direction (*p* = 0.77, paired permutation test), and had a mild statistical increase in the L4 to L2/3 direction (*p* < 0.05, paired permutation test; ***Figure 5—Figure Supplement 2***A). During spontaneous activity, the same type of augmentation did not increase EV relative to the neural-only model in either direction (*p* = 0.095 for L4→L2/3 direction, *p* = 0.93 for L2/3→L4 direction, paired permutation tests; ***Figure 5—Figure Supplement 2***C). Behavior-only models captured a modest yet statistically significant fraction of variance (stimulus: EV=0.02 ± 0.03, spontaneous: EV=0.01 ± 0.01; *p* < 0.001 paired permutation test with shuffled frames). Behavior-only EV correlated with neural EV within areas (stimulus: ~ 0.47–0.50; spontaneous: ~ 0.68–0.69; ***Figure 5—Figure Supplement 2***B,D), indicating that animals/neurons with stronger neural predictability also tended to show stronger behavior-related structure, but in all cases behavior was not driving the main effect.

In monkey neural recordings during resting state condition (eyes open only), pupil diameter provided a small but significant predictive power (EV=0.01 ± 0.01; *p* < 0.001 paired permutation test of prediction vs. shuffled frames prediction). Adding pupil size to the neural model yielded no significant EV gain in either direction (*p* = 1 in the V1→V4 direction, *p* = 0.70 in the V4→V1 direction; ***Figure 5—Figure Supplement 2***E).

Together, these results indicate that behavioral metrics contributed a measurable shared component to cross-area predictability, but they are not the principal driver: most explained variance remains tied to the neural population activity.

### Receptive field overlap and neuronal response properties impact predictability

We investigated which neuronal properties are related to the ability to predict responses by comparing EV and key indicators of neuronal response reliability and receptive field properties, in both visually- and non-visually reliable neurons, during either visual presentation or spontaneous conditions. First, we considered the following properties: (i) max *r*^2^ value (i.e., maximum squared correlation between each neuron in the predictor population and the predicted neuron), (ii) 1-vs-rest *r*^2^ self-consistency (squared correlation between each neuron’s response to one trial repetition of all stimuli/timepoint with the average response of the rest of the trial repetitions; Methods), (iii) SNR of the predicted neuron, (iv) variance across stimuli, (V) variance across repeats of the same stimulus, (vii) variance across timepoints (monkey data only, since we have multiple timepoints per trial), and (viii) split-half *r* (Methods). We plotted EV against each of these variables (mouse: ***Figure 6***B; monkey L: ***Figure 6***E, monkey A: ***Figure 6—Figure Supplement 2***B, monkey D: ***Figure 6—Figure Supplement 3***B) and report the correlation coefficient between EV and each variable in the y-axis in mice (***Figure 6***A) and monkeys (monkey L: ***Figure 6***D, monkey A: ***Figure 6—Figure Supplement 2***A, monkey D: ***Figure 6—Figure Supplement 3***A).

**Figure 6.**
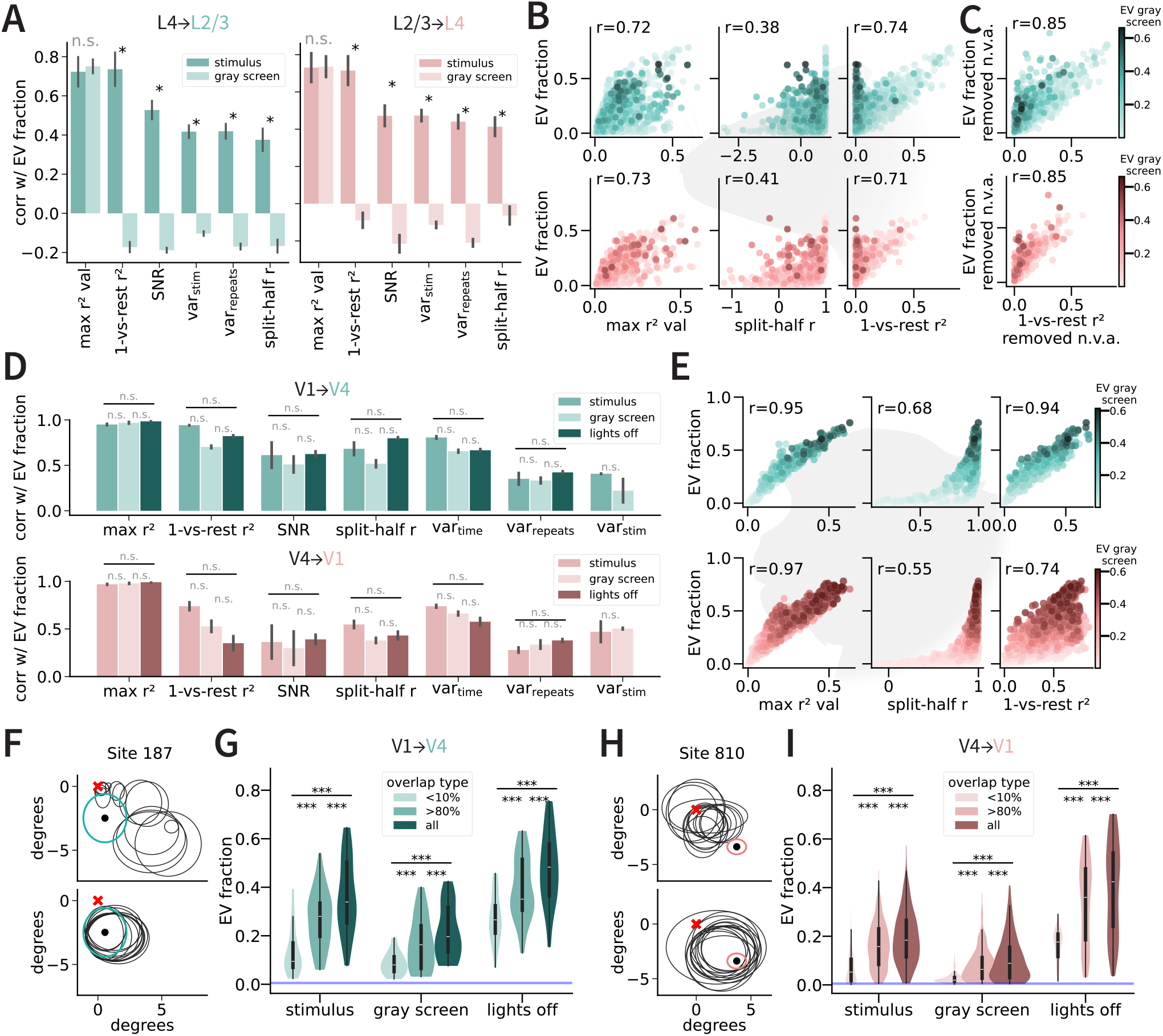
Neuronal predictability depends on SNR, stimulus response variance, and receptive field overlap. **A**. Correlation between different neuronal properties with the predictability of L2/3 (green) and L4 (coral) neuronal responses during the presence (dark color) or absence (light color) of visual stimulus. Neuronal properties measured in mouse V1 include the correlation value of the most correlated pair to each cell (max correlation value, squared), a modified metric of self-consistency (one-vs-rest correlation, squared), SNR, variance in the neuronal activity in response to different stimuli, variance in the neuronal activity across repetitions, and the traditional metric of self-consistency (split-half correlation *r*) (Methods). **B**. Relationship between three neuronal properties and their predictability in a randomly chosen sub-sample of neurons (*n* = 4, 000) for mouse L2/3 (green) and L4 (coral) neuronal responses from both drifting gratings and natural images conditions (combined). Hue represents the degree of predictability for the same neurons during the 30 minutes of gray screen presentation (see color map on bottom right).**C**. 1-vs-rest square correlation relationship with predictability after projecting out dimensions of “non-visual” activity (using gray screen activity (***Stringer et al., 2019a***). **D**. Correlation between different neuronal properties with the predictability of monkey L V4 (green) and V1 (coral) neuronal site recordings during the full-field checkerboard presentation (dark color), gray screen presentation (light color), and lights-off condition (darkest color; solid, hatch lines, and hatch dots). Neuronal properties measured in monkey visual cortex include the max correlation squared value, one-vs-rest squared correlation, SNR, variance across different stimuli (moving bars dataset only), variance across time (within-trial repeat), variance across repeats (within timepoint), and split-half correlation *r*. **E**. Same as B for monkey L V1 and V4 neuronal sites. **F**. Top: Receptive fields of one sample V4 neuronal site (green circle, array 2 electrode 187) and 14 randomly selected V1 neuronal sites as predictors (black circles), constrained on sites that share less than 10% receptive field overlap with the V4 site. Bottom: Receptive fields of the same neuronal site 187 and 14 randomly selected V1 neuronal sites used as predictors, constrained on sites that share at least 80% receptive field overlap with the V4 site. **G**. Differences in predictability of V4 neural activity (*n* = 110 site recordings) in terms of 14 V1 predictor sites with less than 10% RF overlap (light green), 14 predictor sites with at least 80% RF overlap (green), and all predictors (dark green). Predictions were computed for recordings in response to the stimulus presentation (sliding bars and full-field checkerboard images), gray screen presentation, and lights off. **H**. Bottom and top left: Same as F but for monkey L sample V1 site 810. **I**. Same as D, but for V1 (*n* = 970 site recordings).

In mice, during both stimulus presentation and gray screen presentation, the most correlated property with a neuron’s inter-areal predictability was the max *r*^2^ (***Figure 6***A). For the other 5 properties, there was a strong distinction between stimulus presentation (dark bars) and gray screen presentation (light bars): all 5 properties were positively correlated with the neural activity predictability EV fraction during stimulus presentation, but they were slightly anticorrelated with their predictability EV fraction during gray screen presentation. Because the split-half correlation calculation averages out the non-stimulus-dependent variability in both halves of the trials, it showed a weaker correlation with EV, which depends on trial-by-trial modulation. The one-vs-rest *r*^2^ metric, which also examines trial-by-trial modulation and does not average split-half trials, yielded a stronger correlation with EV during stimulus presentation.

When examining the relationship between 1-vs-rest self-consistency and inter-laminar prediction EV in mice, we observed a bimodal distribution of neurons: one group of neurons showed high EV despite having low self-consistency, and another group showed EV correlated with self-consistency (***Figure 6***B third column; ***Figure 6—Figure Supplement 1***A; *p* < 0.001, Hartigan’s dip test after thresholding neurons with *EV* > 0.4). The responses of neurons with low self-consistency also showed high EV during gray screen presentation. This bimodality was present in two of the three mice (*p* < 0.001 for mouse MP031, MP032, *p* = 0.69 for mouse MP033, Hartigan’s dip test; self-consistency and EV fraction relationships across all mice can be seen in ***Figure 6—Figure Supplement 1***B). To better understand the responses of neurons with low self-consistency, we projected out the “non-visual ongoing neuronal activity” from the neuronal responses (***Stringer et al., 2019a***) (Methods). This non-visual ongoing activity is deemed to be influenced by spontaneous behavior (***Stringer et al., 2019b***). Projecting out this non-visual activity eliminated this bimodality (*p* = 0.71 and *p* = 0.98 for L4→L2/3 and L2/3→L4, respectively; Hartigan’s dip test, ***Figure 6***C,D). This observation could be because the responses of neurons with low self-consistency can no longer be predicted, or because the responses of those neurons became more reliable and therefore were highly predicted. To distinguish between these two possibilities, we compared both the 1-vs-rest self-consistency and the prediction EV before and after removing the non-visual activity. Removing the non-visual ongoing activity increased the self-consistency value across the three mice (***Figure 6—Figure Supplement 1***C; *p* < 0.001, paired permutation test). Interestingly, the inter-laminar EV fraction decreased in MP031 and MP032 mice, yet increased in MP033 (***Figure 6—Figure Supplement 1***F*p* < 0.001, paired permutation test). When examining individual pairwise relationships in a fraction of highly predictable neurons (lines in ***Figure 6—Figure Supplement 1***F), we found that some of the highly predictable neurons remained predictable after removing the non-visual activity, whereas other highly predictable neurons dropped their EV fraction dramatically. Together with the small yet reliable shared behavior contributions (***Figure 5—Figure Supplement 2***), these results indicate that a non-trivial portion of inter-areal predictability in mice reflects shared internal-state fluctuations, whereas a complementary portion persists after removing those dimensions, consistent with stimulus-driven interactions.

In monkeys, one of the neuronal properties that showed high correlation with inter-areal prediction EV across all conditions was also the maximum correlation value (first column in ***Figure 6***D, E, ***Figure 6—Figure Supplement 2***A, and ***Figure 6—Figure Supplement 3***). Other neuron properties like SNR, split-half correlation, and one-vs-rest correlation were also highly correlated with inter-cortical predictability EV(***Figure 6***D, ***Figure 6—Figure Supplement 2***A, and ***Figure 6—Figure Supplement 3***A). The split-half correlation was highly correlated with EV fraction, although the relationship was highly non-linear (middle column in ***Figure 6***D, ***Figure 6—Figure Supplement 2***B, and ***Figure 6—Figure Supplement 3***B). Using the one-vs-rest squared correlation removed some of this non-linearity and further increased the correlation with the EV fraction in monkeys L and A (third column in ***Figure 6***D, ***Figure 6—Figure Supplement 2***B). In addition, the relationship between one-vs-rest correlation squared and EV fraction did not show evidence of bimodality (*p* = 0.99, Hartigan’s dip test focusing on sites with *EV* > 0.4).

We conjectured that neurons that have overlapping receptive fields (RFs) should share more information, and therefore their responses would better predict each other than neurons with non-overlapping RFs. In addition, even when all neurons are exposed to the same stimulus (full field symmetrical checkerboard image, gray screen, darkness, etc), neurons with overlapping RFs may be more synaptically connected, resulting in better inter-cortical predictions. To test this hypothesis, we compared inter-cortical predictions in different ensembles of neurons with RFs that differed in their degree of overlap. This hypothesis was only tested in monkeys L and A because we did not have access to RF estimates in the mouse data, and we did not have enough sites with RF overlap in monkey D. For each V4 site whose responses we predicted, we separated the predictors into two groups with matched size: one group where all the V1 predictor sites had <10% RF overlap (sample of one V4 site in monkey L, ***Figure 6***F, top), and one group where all the V1 predictor sites had >80% RF overlap (sample of one monkey L V4 site, ***Figure 6***F, bottom). A similar procedure was followed when predicting the activity of V1 neurons from V4 predictor neurons (***Figure 6***H). Inter-areal prediction was higher in the >80% RF overlap condition compared to the <10% RF overlap ensembles in both directions and across all stimulus conditions (***Figure 6***G, I for monkey L; ***Figure 6—Figure Supplement 2***C, D for monkey A). To compare these numbers with those obtained with larger numbers of sites, we calculated the EV performance while using all predictors. In most cases, predictions using the >80% RF overlap group were lower than when using all predictors, showing that predictor neurons with low RF overlap still contributed to the EV (***Figure 6***G,I, for monkey L; ***Figure 6—Figure Supplement 2***C,D for monkey A).

### Inter-areal predictability is both stimulus and non-stimulus driven

The results in ***Figure 5*** and ***Figure 6*** pointed to components of the predictable responses that are stimulus-driven and other components that are not stimulus-driven. We considered two alternative hypotheses: (1) If shared information were strictly stimulus-driven, prediction across repeats of the same stimulus should remain accurate; (2) If shared information were entirely stimulus-independent, prediction across repeats should approach chance (*EV* ≈ 0). To test these hypotheses, we compared the inter-areal prediction EV fractions using unshuffled versus shuffled trials. We tested these hypotheses within each species, and our goal was not to compare the two. Shuffling was performed across repeat trials of the same images (mice: ***Figure 7***A, monkeys: ***Figure 7***D). In mice, one stimulus presentation was either a drifting grating or a natural static image. In monkeys, one stimulus presentation was either the one checkerboard image, a large, thick, fast-moving bar, or a small, thin, slow-moving bar. In both species and in both directions, inter-areal prediction EV fraction decreased after shuffling stimulus repeats compared to before shuffling (***Figure 7***B for mouse; ***Figure 7***E for monkey L; ***Figure 7—Figure Supplement 1***A for monkey A; ***Figure 7—Figure Supplement 1***C for monkey D; *p* < 0.001 for both species and directions, paired permutation test).

**Figure 7.**
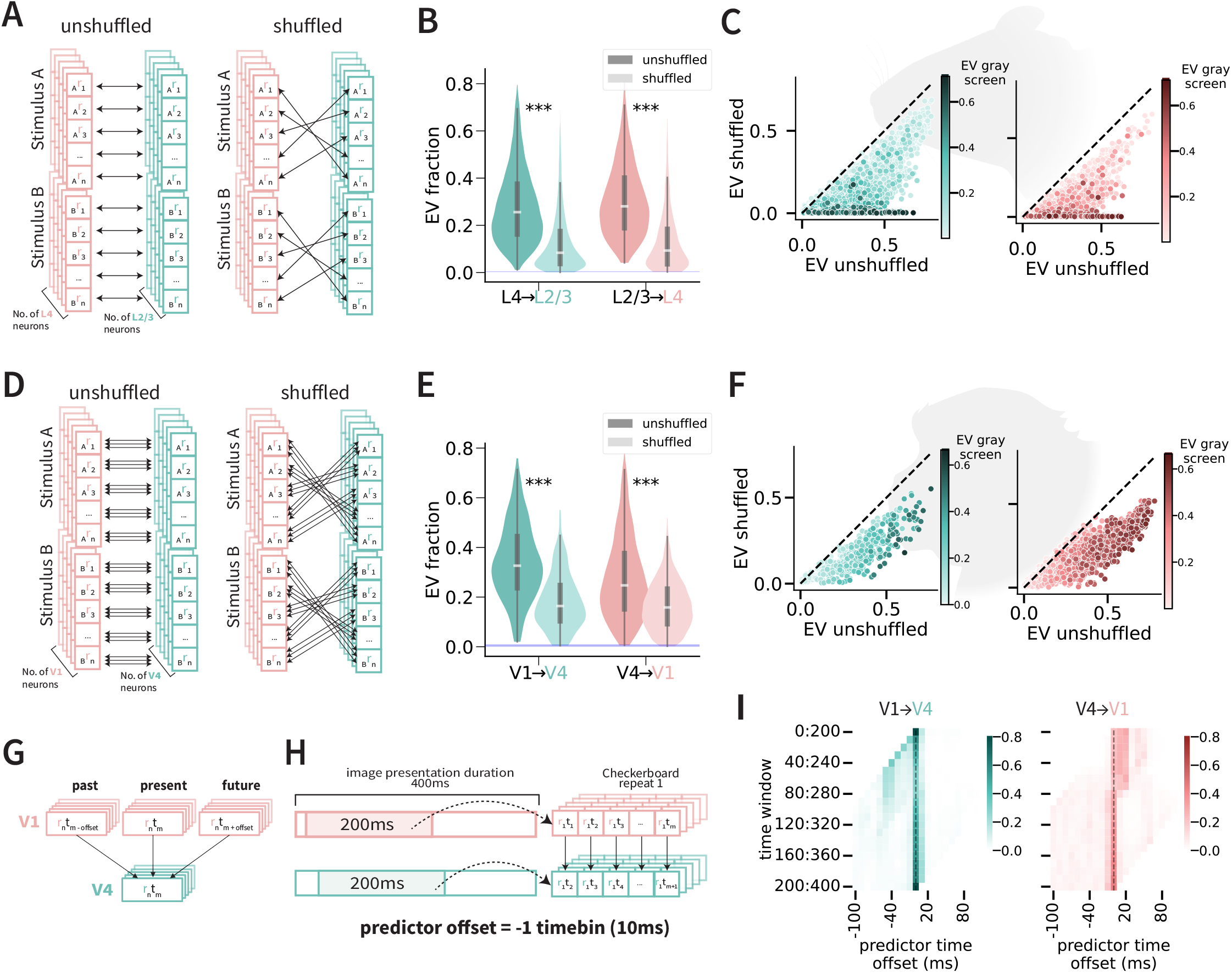
Predicting neuronal activity across time reveals shared stimulus- and non-stimulus driven information in both mouse and monkey visual cortex, along with latency and non-latency specific correlations in monkey V1/V4. **A**. Shuffled-trial-repeat experiment set-up for comparing unshuffled vs. shuffled prediction in mouse V1 L2/3 and L4. Neuronal activity in response to stimulus repeats was shuffled within their respective image. **B**. EV fraction for unshuffled (dark) and shuffled (light) trials in the L4 → L2/3 (green) and L2/3 → L4 (coral) directions. **C**. Relationship between shuffled (y-axis) and unshuffled (x-axis) trial repeat EVs in the mouse L4 → L2/3 (left, green) and L2/3 → L4 (right, coral) directions. Hue represents EV fraction during gray screen activity. **D**. Shuffled-trial-repeat experiment set-up comparing unshuffled vs. shuffled prediction in monkey data. Neuronal activity (including all timepoints) in response to stimulus repeats was shuffled within their respective image. Since the checkerboard presentation was only one stimulus, the experiment visualization only applies to the “Stimulus A” portion. **E**. Same as **B** for monkey L V1 → V4 (green) and V4 → V1 (coral). **F**. Same as **C** for monkey V1 → V4 (green) and V4 → V1 (coral). **G**. Illustration of time offsets applied to monkey neuronal activity for inter-areal predictions. Instead of neuronal activity prediction between areas being done on simultaneous activity (middle coral and bottom green box), the V4 neuronal activity (green) at time *t*_*m*_ was predicted using V1 neuronal activity (coral) at time *t*_*m*±*offset*_, were *offset* represents 1–8 timebins (25 ms per timebin) before (if negative; left coral box) or after (if positive; right coral box) time *t*_*m*_. Time offset experiment was done in both prediction directions (V1 → V4 and V4 → V1). **H**. Experimental set-up example for predicting neuronal activity in V4 using V1 neuronal activity from 10 ms prior to V4 activity. Neural activity is in response to a repeated checkerboard image. A 200 ms section of the cortical area was used to represent the image presentation response, and was offset −1 timebin (10 ms) to predict a 200 ms target cortical area. During the prediction experiments, the 200 ms window was slid across the entire duration of the stimulus **I**. Time offset prediction results across both V1→V4 (left, green) and V4→V1 (right, coral) prediction directions. Each square corresponds to the fraction of neuronal sites whose neural activity were best predicted during that offset period and time window.

Interestingly, the EV fraction during shuffled trial-repeats still remained significantly above chance (*p* < 0.001, paired permutation test of prediction vs. shuffled frames prediction). In mice, neurons showed two types of distribution in terms of their response predictability in shuffled and unshuffled trials. For a subset of neurons, the EV fraction was still high in the shuffled condition, albeit their EV was lower than in the unshuffled case (***Figure 7***C; points below but near the diagonal line). For another subset of neurons, the EV fraction during shuffled trials was much lower or even near zero. The responses of the latter group had the highest predictability during gray screen activity.

In monkeys, neurons farther away from the diagonal line showed higher EV fraction during gray screen activity (***Figure 7***F for monkey L; ***Figure 7—Figure Supplement 1***F for monkey A; ***Figure 7—Figure Supplement 1***G for monkey D). In addition, we examined whether shuffling repeat presentations of gray screen images (simulating spontaneous activity) would result in any prediction at all. There was a more profound decrease in inter-cortical performance in all monkeys (***Figure 7—Figure Supplement 2***A–C) with no neurons that remained as predictable during shuffled trials compared to unshuffled trials (***Figure 7—Figure Supplement 2***E–G).

### Accounting for latency differences improves inter-areal activity predictions in monkeys visual area sub-populations

Given the latency differences in neuronal responses between V1 and V4 ***Schmolesky et al., 1998***, we asked whether accounting for this latency could result in better inter-cortical prediction. To test this hypothesis, we offset the neuronal activity using different lags for each area (***Figure 7***G, H) and recalculated the ridge regression predictions. For each offset level, we calculated the percentage of neurons where the EV fraction peaked at that offset. For the checkerboard image, in monkeys L and A V1→V4 predictions, the biggest percentage of neurons had a peak performance when there was no time offset between areas, with a substantial proportion of neurons with a peak performance for 10–30 ms offsets in the negative direction (i.e., V1 activity preceding V4 activity; ***Figure 7***I, left for monkey L, ***Figure 7—Figure Supplement 1***E, left for monkey A). This distribution of peak EV values was only present during early visual responses (first 275 ms of stimulus onset). In monkeys L and A V4→V1 direction, there was a large proportion of neurons with peak EV when considering 10–20 ms offsets in the positive direction (i.e., V4 after V1, ***Figure 7***I, right for monkey L, ***Figure 7—Figure Supplement 1***E, right for monkey A). These differences were apparent in the early part of the visual response, before 250 ms. When offsetting the neuronal responses to gray screen presentations, across all times and areas, the highest percentage of neurons with peak EV was when there was no time offset (***Figure 7—Figure Supplement 2***D).

### Neural signals can also be predicted at the ensemble level using local field potential signals

We repeated the analyses in monkeys L and A using the local field potential (LFP) signals. We evaluated whether the predictability of LFP signals across areas depended on the signal frequency by considering band-limited LFP amplitudes (Low: 2–12 Hz; Beta: 12–30 Hz; Gamma: 30–45 Hz; High-gamma: 55–95 Hz; Hilbert envelopes, z-scored). In both monkeys, the Gamma band showed a feed-forward signature in the early evoked period: the V1→V4 predictability peaked at negative offsets (~10–30 ms; V1 leads), and the V4→V1 predictability peaked at positive offsets of similar magnitude ***Figure 7—Figure Supplement 3***A,D for monkey L; ***Figure 7—Figure Supplement 3***B,E for monkey A), consistent with previous findings (***Semedo et al., 2022; van Kerkoerle et al., 2014; Bastos et al., 2015***). The Low and Beta frequency bands exhibited a broader structure with less consistent peaks in predictability offset timing.

## Discussion

Neuronal activity in one brain region or layer within the visual cortex can be used to predict neuronal activity in another nearby and anatomically connected region or layer in single trials (***Figure 2***). In monkeys, predictability was asymmetric: V1 activity better accounted for V4 activity than vice versa (***Figure 3, Figure 7***). This inter-areal prediction persisted across different stimuli (***Figure 4***) but also in the absence of a visual stimulus, during gray-screen and lights-off periods (***Figure 5***). The degree of predictability increased with signal-to-noise ratio, response variance, and the degree of overlap between receptive fields (***Figure 6***).

In line with other studies in mice (***Stringer et al., 2019b; Niell and Stryker, 2008; Andermann et al., 2011***), we observed an approximately bimodal distribution of neuronal responses, with a large subset of neurons that do not show reliable responses to visual stimuli both in L4 and L2/3 (***Figure 6***B, third column; ***Figure 6—Figure Supplement 1***A). However, even for the subset of neurons considered to be “non-visual”, at least within the set of stimuli and conditions examined here, their activity remains predictable. This bimodal distribution disappears when projecting out potential non-sensory ongoing activity (***Figure 6***C; ***Figure 6—Figure Supplement 1***D) (***Stringer et al., 2019b, 2021***). At the population level, neuronal encoding subspaces in mouse visual cortex have been shown to have little overlap between visual sensory and non-sensory (behavioral) information, with only one shared dimension (***Stringer et al., 2019b***). The visually unreliable, yet highly predictable, subset of neurons described here could be the neuronal group driving this orthogonality. As expected, the activity of “visual” neurons can be better predicted during visual presentation and not during gray screen presentation. In stark contrast, the activity of non-visual neurons can be predicted even better during gray screen presentation than during visual stimulation. There was no such bimodal distribution in the data from monkeys. One possibility is that there may be no (or very few) non-visual neurons in monkey V1 or V4. Indeed, the overwhelming majority of recording sites in V1 and V4 responded strongly to visual stimulation. Yet, the comparisons between the results in mice and monkeys reported here need to be interpreted with caution because the two datasets differ in terms of recording techniques (electrophysiology versus two-photon imaging), consequently also in their temporal resolution (one millisecond versus hundreds of milliseconds), and the type of interaction studied (across areas versus across layers), in addition to any differences between species.

In monkeys, sites where the receptive fields (RF) of V1 and V4 overlap can better predict each other compared to other sites showing little RF overlap. This observation could reflect RF-dependent intrinsic connectivity between areas, but also RF-dependent shared inputs from other areas, such as the thalamus. In the latter case, those putative shared inputs cannot be strictly dependent on visual inputs, given that the effect of RF overlap persists even during gray screen conditions.

Many computational models that aim to explain neuronal activity in the visual cortex are based on feedforward signal propagation, with increased receptive field sizes, selectivity, and feature invariance along the visual hierarchy (***Serre et al., 2007b; Kreiman, 2021; Connor et al., 2007; Kreiman and Kriegeskorte, 2012***). Consistent with this idea, we described an asymmetry in the degree of predictability, with V1 neurons explaining V4 responses better than the other way around. This observation persisted after controlling for neuronal count and split-half correlation values and was also apparent during the lights-off condition. In contrast, there was no asymmetry when comparing inter-laminar prediction directions in mice. The lack of asymmetry in inter-laminar prediction directions in mice could be due to the slow dynamics in calcium imaging, the lack of a clear inter-areal hierarchy, or differences between species.

The asymmetry in directionality was also observed when implementing temporal delays to inter-areal prediction, consistent with processing delays across areas (***Semedo et al., 2022; Gokcen et al., 2022; Schmolesky et al., 1998***). A substantial proportion of neurons increased their inter-areal predictability when offsetting the times between areas, specifically in the direction that aligns their neuronal activities. In contrast to the temporal delays associated with processing visual stimulation, during gray screen presentation, the activity of most neurons was best predicted in the absence of any time offsets, suggesting that the internally generated neuronal activity during spontaneous conditions may be largely driven in a non-feedforward manner. Further evidence supporting the distinction between visually-driven and non-visually-driven interactions comes from the observation that trial repeat shuffling reduced, but did not eliminate, predictability in both mice and monkeys. In mice, when comparing shuffled versus unshuffled activity, we encountered again a bimodal distribution, where a group of neurons was closer to the diagonal line (their responses were predicted as well during the shuffled compared to the non-shuffled condition), and another group of neurons that were closer to the x-axis (their responses could not be predicted during the shuffled condition). The responses of the latter group were best predicted during gray screen activity, suggesting that they mostly shared non-visual information. The predictive power in mouse V1 from layer 4 to layer 2/3 during spontaneous conditions has been recently shown in ***Papadopouli et al. (2024)***, consistent with our findings. The overall area population decrease in predictability after shuffling may be due to the influence of non-visual activity, such as movement (***Stringer et al., 2019b***), especially since these non-visual stimulus effects have been shown to occur on a timescale of approximately one second, consistent with the results in our study. In macaques, context-dependent effects are likely not due to movement, since the monkeys maintained fixation during the stimulus task, and visually-evoked activity was not driven by movement (***Talluri et al., 2023***).

The neural recordings in mice and monkeys are different in terms of: (i) recording modalities (calcium versus electrophysiology), (ii) temporal resolution (hundreds of milliseconds versus milliseconds), (iii) neuronal types, (iv) SNR, (v) cortical targets (layers versus areas), (vi) sample sizes, (vii) stimuli, and (viii) task conditions (***Stringer, 2018; Chen et al., 2022b***). The goal of this study was not to make any direct quantitative comparison across species, but rather to investigate the properties of inter-areal interactions within each species.

Several limitations in this study are worth considering in future work. First, neither calcium imaging nor MUAe recordings capture the timing of single spikes in individual neurons. One of many ways in which the timing of individual spikes could be important is that synchronous or close-to-synchronous inputs can have a larger impact on post-synaptic responses (***Salinas and Sejnowski, 2001***). Second, this study focused on linear predictability, but other non-linear models could better capture neuronal activity. Third, and critically, the experimental data evaluated here provide only a fraction of the inputs to a given neuron. For example, the predictor signals studied here are likely to exclude most (in monkeys) if not all (in mice) inhibitory inputs (***Jiang et al., 2015; Shen et al., 2020; Ibrahim et al., 2020; Schuman et al., 2021***). Any cortical neuron receives extensive local inputs and non-local inputs from many other areas. Fourth, we only had limited access to non-visual sources that could contribute to predicting neuronal responses; other sources of input could include finer behavioral measurements, attention, familiarity, and arousal state. Finally, biophysically realistic models of the transformation between inputs and outputs of a given neuron should include their dendritic locations and specific synaptic potentials (***Park et al., 2019; Petousakis et al., 2023***). Therefore, the quantitative results on the prediction of neuronal responses reported here constitute a lower bound.

We evaluated inter-areal interactions in different types of neuronal recordings, timescales, and species. These interactions were assessed in single trials, separating visually-driven and non-visual contributions, and accounting for the directionality and dynamics of neuronal responses. These efforts constitute an initial step towards, and a lower bound for, systematically building computational models that can account for the transformations from sensory inputs to their encoding across multiple processing steps in the cortex. Whereas many efforts with artificial neural networks merely map inputs to outputs (***Kreiman, 2021***), a detailed and systematic understanding of individual processing steps can lead to better and biologically more relevant computational models of sensory processing.

## Methods

### Datasets

Animal characteristics, including sex, age, and genotype/strain, for the previously published mouse and macaque datasets are reported in ***Stringer et al. (2019a***,b) and ***Chen et al. (2022b***), respectively. We used the mouse dataset from (***Stringer et al., 2019a***) containing calcium-imaging activity measurements from 43,630 neurons in layer 4 (L4) and 12,060 neurons in layers 2/3 (L2/3) in V1 of 4 mice (*Mus musculus*; transgenic lines expressing GCaMP6s in excitatory cortical neurons; full strain, sex and age details are reported in ***Stringer et al., 2019a***,b) during 32 different randomly interleaved presentations of either drifting gratings or gray-scale natural images (each one repeated more than 90 times), along with spontaneous activity during 30 minutes of exposure to a gray/black screen (***Figure 1***A, data acquisition details in ***Stringer et al. (2019a***,b)). Calcium imaging activity was recorded during stimulus presentations at a scan rate of 2.5 Hz or 3 Hz (each frame was acquired every 400 ms or 333 ms). The computed stimulus responses per stimulus presentation were averaged based on two frames immediately post stimulus onset. Cortical layers were determined using the 10-12 planar z-positions retrieved during the multi-plane calcium activity acquisition. For stimulus-response and spontaneous recordings, neuronal activity of each neuron was z-scored using its 30-minute gray screen spontaneous activity (mean gray-screen activity subtracted and divided by gray-screen activity standard deviation).

We used the dataset from ***Chen et al. (2022b***) for monkeys A and L and conducted new neural recordings from one additional monkey (monkey D). The Chen et al. dataset consists of envelope multiunit activity (MUAe) and LFP from 1,024 recording sites in one monkey (monkey L) in response to either multiple-day recordings of more than 60 repetitions of a full-size checkerboard image, moving small-thin or large-thick bars in 4 directions, gray screen presentations, or more than 30 minutes of baseline activity where the monkey was in a room with the lights off (***Figure 1***B). We also used data from a second monkey (monkey A) from the same published dataset. However, due to the degraded state of the electrodes at the time of the recording, we were only able to obtain enough data from V4 sites for responses to the full-size checkerboard image, moving bars, and gray screen presentations. Neuronal activity was summed over 25 ms non-overlapping bins. Activity duration was 300 ms, 400 ms, and 1 s for gray screen, checkerboard, and moving bar presentations, respectively. For the recordings during visual stimulation, the neuronal activity was normalized by subtracting the mean activity during the gray screen presentations separately for each site. We collected new data from an additional monkey in response to the full-size checkerboard image presentations, gray screen presentations, and lights-off activity (~15 minutes). Monkey D (male rhesus macaque, *Macaca mulatta*, 12 years old, 14.4 kg) was chronically implanted with floating multielectrode arrays (Microprobes for Life Science, Gaithersburg, MD, USA) targeting V1, V4 and additional visual areas. The present study analysed the V1 and V4 arrays only (16 channels per area; 32 electrodes). All procedures received ethical approval by the Harvard Medical School Institutional Animal Care and Use Committee (IACUC protocol IS0003351-3) and conformed to NIH guidelines provided in the Guide for the Care and Use of Laboratory Animals. All relevant ethical regulations for animal and nonhuman primate testing and research were followed.

The same conditions of stimulus presentation (image/square size, presentation time, etc) were presented as in ***Chen et al. (2022b***). Stimulus presentation and data acquisition (including electrophysiology and eye tracking) were coordinated using MonkeyLogic2 software (Hwang et al., 2019) and OmniPlex Neural Recording Data Acquisition Systems (Plexon Inc., Dallas, TX, USA), integrated via custom MATLAB scripts. Online spike sorting was performed with the PlexControl client using waveform-based classification. Visual stimuli were displayed on ViewPixx EEG monitors (VPixx Technologies, Saint-Bruno, QC, Canada; 1920×1080, 120 Hz), and eye position was monitored via ISCAN (ISCAN Inc., Woburn, MA, USA). Raw analog signals were used to calculate MUAe using same processing logic as ***Chen et al. (2022b***).

### Visual Reliability

A neuron or site was defined to be visually reliable if its signal-to-noise ratio (SNR) was 2 or higher and its split-half correlation value was 0.8 or higher. Due to the high number of repetitions of visual stimuli, the split-half correlation was skewed toward high values, which is why we used a higher split-half correlation threshold than commonly used in other studies. In mice, the SNR for each neuron was calculated as:

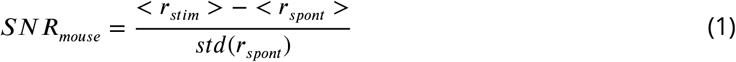

where <> denotes mean, std denotes the standard deviation, *r*_*stim*_ is the average activity in response to stimuli, and *r*_*spont*_ indicates the average activity over the 30-minute gray screen presentation.

In monkeys, we followed the definition in ***Chen et al. (2022b***) and calculated the SNR for each site as:

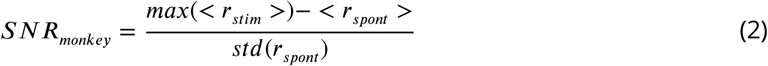

using the peak activity during the checkerboard presentation for the signal, and the average gray screen neuronal activity as background (denoted as *r*_*spont*_).

In mice, the split-half consistency was calculated by correlating the average activity for the 32 stimuli in half of the trials, randomly chosen, with the average activity in the other half of the trials, followed by Spearman-Brown correction (used to correct for the division of trials by half). In monkeys, during checkerboard presentations, the split-half consistency was calculated by correlating the average activity of the 16 timepoints (0–400 ms; 25 ms bins) of checkerboard presentation of 25 random trial repetitions with the average activity of another non-overlapping 25 random repetitions, followed by Spearman-Brown correction. During moving bar presentations, the 40 time-points (0–1s; 25 ms bins) during 25 random trial repetitions were first concatenated across the 4 directions (total of 160 timepoints), and then correlated to the concatenated averaged activity of another nonoverlapping 25 random trial repetitions, followed by Spearman-Brown correction. For all split-half consistency calculations, we randomly sampled trials 20 times.

### Inter-areal predictions

Let 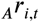 be the activity of neuron or site *i* in area *A* at timepoint *t*, where *A* can be L4 or L2/3 in the mouse data and V1 or V4 in the monkey data. Neuronal activity from one area (e.g., mouse V1 L4 or monkey V1) was used to predict activity in the other area (e.g., mouse V1 L2/3 or monkey V4) using ridge regression (***Figure 1***E). The activity of each neuron *i* in area A2 was predicted from *n*_*A*1_ neurons in area A1 as follows:

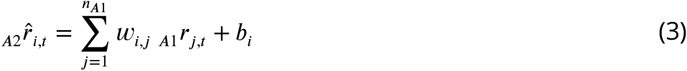

During fitting, we minimized the residual sum of squares (RSS), defined as:

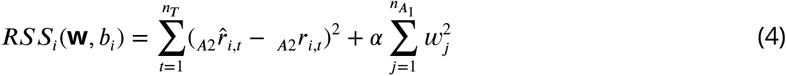

where **w** is the weight vector for predicting the activity of neuron *i, n*_T_ is the number of images/time points and *α* controls the regularization strength (*α* was tuned for each dataset with an independent sample and ranged from 10^3^ to 10^5^). Predictability for each neuron was evaluated using 10-fold cross-validation across trials and quantified as squared Pearson’s r between held-out _*A*2_*r*_*i,t*_ and 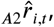, referred to as explained variance fraction (EV fraction) throughout.

To remove temporal auto-correlation that would inflate the apparent prediction despite cross-validation, we removed training timepoints near the test timepoints closer than the decay window of the activity auto-correlation (mouse: 5 s; monkey: 1.5 s). The auto-correlation decay window was determined using time-series forecasting Ridge Regression (using *r*_*t*_ to predict *r*_*t*+*d*_, where *d* represents a delay). The delay was increased until the EV fraction approached chance.

To model a Poisson distribution of MUAe responses, we also fit a Poisson generalized linear model (GLM) using the same predictors as above. We implemented this model with scikit-learn’s `PoissonRegressor` (log link; intercept *b*_*i*_ unpenalized) and tuned *α* per dataset using an independent sample. Model evaluation followed the same 10-fold cross-validation and temporal exclusion procedure as ridge regression (1.5 s exclusion window for monkey). Predictability was quantified as the squared Pearson correlation between observed and predicted MUAe on held-out folds (EV fraction), allowing direct comparison between ridge regression and Poisson GLM (***Figure 2—Figure Supplement 1***M).

### Prediction directionality

We compared predictability across layers in different directions (in mice: L4→L2/3 vs. L2/3→L4) and also predictability across areas in different directions (in monkeys: V1→V4 vs. V4→V1) (***Figure 3***). To ensure that results were not dependent on the number of neurons/sites, we randomly subsampled the number of neurons/sites of the area containing the larger number of neurons/sites (L2/3 for mouse; V1 for monkeys) to match the number of predictors in both directions (10 iterations per recording type, neuron count details in ***Table 1*** and ***Table 2***. Results in ***Figure 3*** represent the median across the 10 subsamples). To account for potential changes in intrinsic predictability, we ensured that the neurons from both areas were matched in terms of the distribution of split-half correlation values so that the difference between individual area neurons/sites was less than 0.002. To assess the intrinsic predictability of neurons/sites in each region, the areas were used to predict themselves, where one neuron/site in the area was predicted by the remaining neurons in the same area. This “intra-areal” prediction was used to normalize EV fraction to compare directionality of prediction.

### Stimulus types and spontaneous activity comparison

We compared predictability for different stimulus conditions (***Figure 4, Figure 5***). To compare inter-areal prediction across stimulus types and between the presence or absence of stimuli, the number of predictors (neurons or sites) and timepoints was sub-sampled to be the same across all datasets. In monkeys, the time spent recording the lights-off condition was much greater than during stimulus or gray screen presentations. To account for the difference in time duration and therefore training size, we subsampled time periods to be the same across all stimulus, gray-screen, and lights-off, lights-off eyes open, and lights-off eyes closed conditions.

### Neuron properties

We compared different neuronal properties with predictability measurements (***Figure 6***). The SNR and split-half correlation has been defined above. The absolute max pairwise correlation value of each neuron/site *i* in one area with all neurons in the other area was calculated as

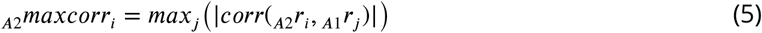

where _*A*2_*r*_*i*_ represents the activity of neuron/site *i* in area A2, which are correlated with the activity of every neuron *j* in area A1 (denoted as _*A*1_*r*_*j*_).

One–vs–rest self-consistency (1-vs-rest *r*^2^). For each neuron/site *i* with _T_ repeat trials, let 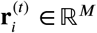 be its activity vector from repetition *t* across all stimuli/timepoints (length *M*, defined below). The leave-one-out mean response over repetitions *excluding t* is

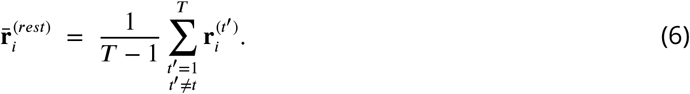

Throughout, the superscript (*rest*) means “all repetitions excluding repeat *t*”.

The one–vs–rest correlation for the held-out repetition *t* is

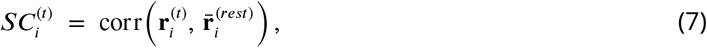

We then average these correlations across all held-out repetitions:

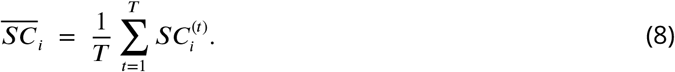

In mice, 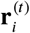 stacks the responses to the *S* = 32 stimuli for repetition *t* (each element is the mean of the two frames immediately post-onset). In monkeys, during checkerboard presentations, 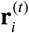 stacks the _T_ = 16 timepoints (0–400 ms; 25 ms bins). In monkey moving-bar presentations, 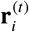 concatenates the _T_ = 160 timepoints (4 directions × 40 bins; 0–1 s, 25 ms bins). Variance metrics. All variances were computed on gray–baseline–subtracted signals. In mice (32 gratings, *K* repeats), for neuron *i*, let 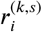 denote the scalar response to stimulus *s* ∈ {1, …, *S*} on repeat *k* ∈ {1, …, *K*} (repeat-averaged within the analysis window). We defined:

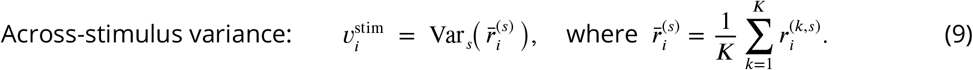

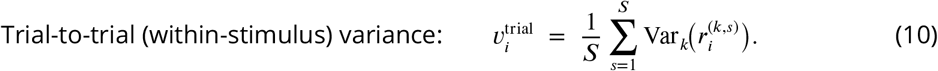

In monkeys, for the checkerboard stimuli, for site *i* let 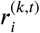 denote the scalar response to time-point *t* ∈ {1, …, _T_} on repeat *k* ∈ {1, …, *K*}

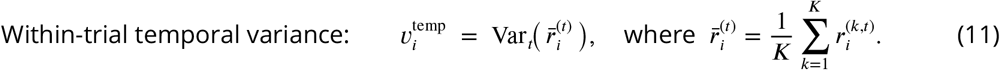

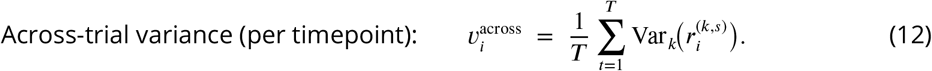

In monkeys, for the moving bar stimuli, the same two metrics were computed per stimulus *s* and then averaged across the four stimuli. In addition, we defined an across-stimuli variance of the time-averaged, trial-mean response:

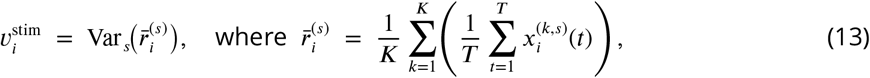

These variance covariates (together with SNR, split-half self-consistency, and one-vs-rest *r*^2^, where applicable) were included as regressors in the *within–dataset-type* linear models used to obtain residual explained variance, 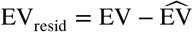.

### EV residualization

For each direction (V1→V4, V4→V1) and stimulus, we fit an ordinary least squares (OLS) model:

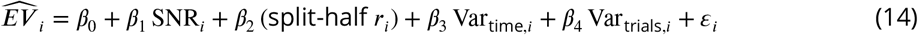

where 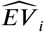 is the predicted explained variance of each neuron/site *i*. Predictors were z-scored within each direction or stimulus.parisons (***Figure 3—Figure Supplement 2***E–H; ***Figure 4—Figure Supplement 1***K,O). SNR_*i*_ is its signal-to-noise ratio (defined in the *Visual Reliability* section), split-half *r*_*i*_ is its split-half consistency (also defined above), Var_time,*i*_ and Var_trials,*i*_ are the temporal and across-trial variance metrics (defined in the *Variance metrics* section), β_0_, …, β_4_ are regression coefficients, and ε_*i*_ is the residual error term. Predictors were z-scored within each direction or stimulus. Residual EV was defined as 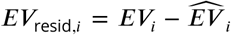 and used for all post-adjustment comparisons (***Figure 3—Figure Supplement 2***E–H; ***Figure 4—Figure Supplement 1***K,O).

### Dip test of unimodality

We used Hartigan’s dip test to evaluate whether the distribution of 1-vs-rest squared correlation values was unimodal or not (***Figure 6—Figure Supplement 1***A,D). The test was performed after restricting the analysis to highly predictable neurons/sites (*EV* > 0.4). The dip statistic measures the maximum deviation between the empirical cumulative distribution function and the closest unimodal distribution (***Hartigan and Hartigan, 1985***). Reported *p*-values correspond to the probability of obtaining a dip statistic at least as large under the null hypothesis of unimodality. Dip tests were computed separately for each cortical layer and mouse (Python package`diptest`).

### Receptive field overlap comparisons

In monkeys L and A, receptive field (RF) ellipses were calculated using center and edge locations in the dataset (***Figure 6***E, F). To calculate the percentage of RF overlap between the neuronal sites to be predicted and the predictor, the intersection area between ellipses was retrieved using the Shapely Python package, and divided by the area of the predicted site. Sites that had predictors that overlap both more than 80% and less than 10% were selected to compare inter-areal predictions. To control for predictor size, 14 random predictor sites from all the sites in each overlap type were subsampled (10 random samples without replacement).

### Trial repeat shuffling and time offset predictions

For the shuffled-trial experiments, we shuffled the predictor activity across repeat trials showing the same stimulus (***Figure 7***). Thus, the stimulus order remained the same. For the mouse time-offset analysis, the activity of predictor neurons was time-shifted in the positive or negative direction, with 1 bin corresponding to 1 stimulus presentation (800–900ms). For the monkey dataset, the predictor activity (400 or 1000 ms per presentation, 16–50 bins of 10 ms each) was offset across time bins. We used sub-windows of 200 ms to avoid window-length differences that would otherwise be introduced if we shifted the entire trial response.

## Data and code availability

All the computational models and code for data analyses are publicly available at this link. All the data are publicly available for mouse (***Stringer et al., 2019a, 2018***), for monkeys L and A (***Chen et al., 2022b***,a), and and for monkey D (***Hidalgo, 2026***). The newly generated monkey D dataset is released under a CC BY 4.0 licence, and the analysis code is released under the MIT license.

## Author contributions

The experiments and tasks were designed by DH, GD, WX and GK, with extensive discussion and help from MP and SS. All the data analyses were performed by DH and GD. The new macaque neurophysiology data were collected by GRT and CP. The manuscript was written by DH and GK, with input from all authors.

## Acknowledgments

The authors would like to thank Elisa Pavarino, Leonardo Polina, Sara Djambazovksa, Jan Drugowitsch, Wei-Chung Allen Lee, Morgan Talbot, for providing comments on the manuscript. Figures were partially created in BioRender. Hidalgo, D. (2026) https://BioRender.com/voyf0rq.

---

**Figure 2—figure supplement 1.**
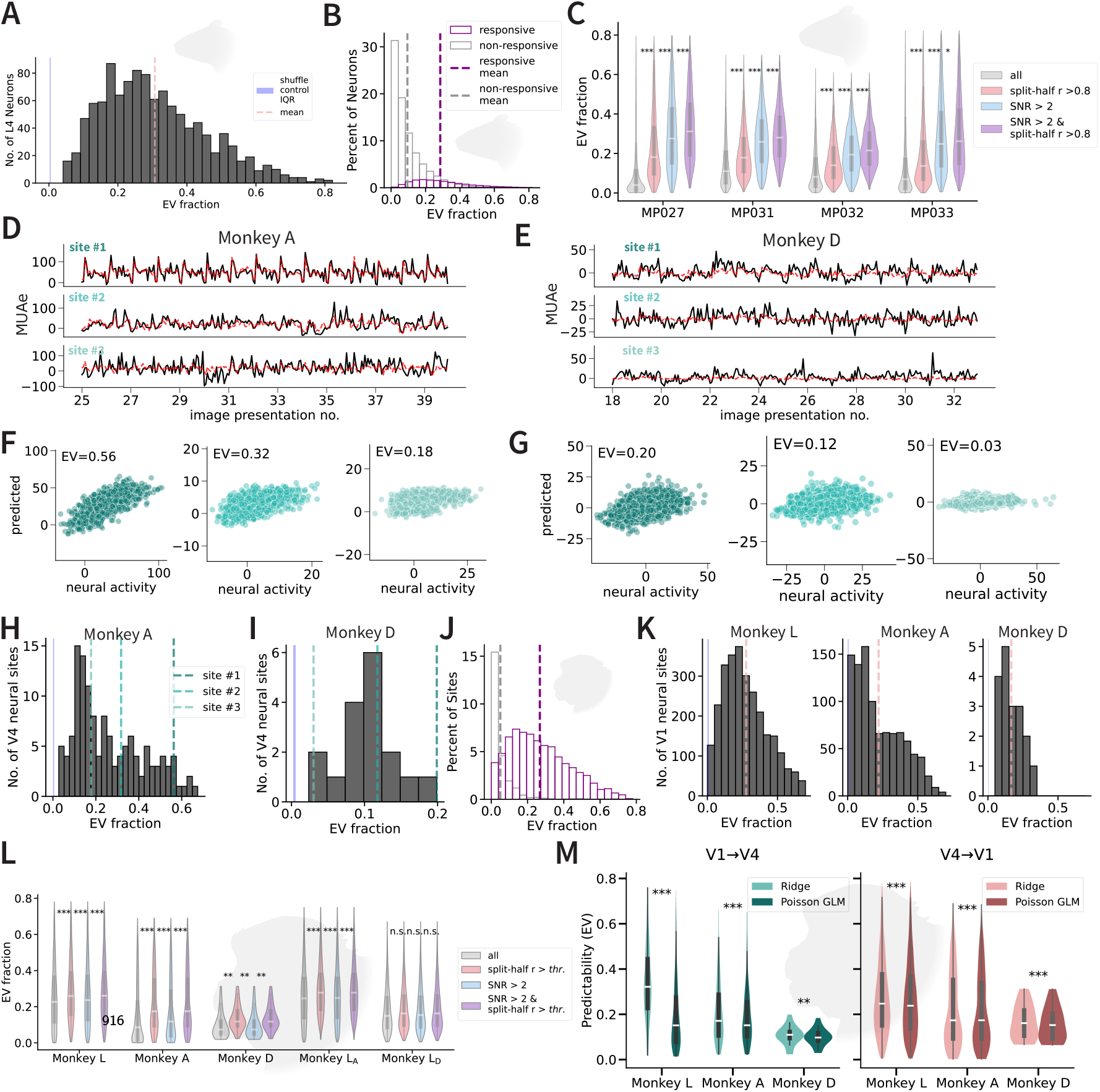
EV fraction in mouse L4 neurons and monkey V1 neuronal sites and comparison between visual and non-visual neurons/sites. **A**. Distribution of EV fraction in L2/3→L4 regressions in cells deemed visually reliable in 4 mice and 7 recording days (*n* = 1113 neurons). **B**. Distribution of EV fraction values for visually (purple) and non-visually (gray) reliable neurons in mouse L2/3 and L4. In mice, we used a conservative criterion to select neurons that were visually reliable, based on an average signal-to-noise ratio over 2 and a split-half correlation value of at least 0.8 (Methods). **C**. Differences in EV fraction using different filtering methods to determine visually reliable neurons in mouse L2/3 and L4 across the 4 mice. **D**. Example of monkey A MUAe activity (black) in response to a full-field checkerboard image in three V4 neuronal sites along with regression-model predictions (red) for a typical site (2, middle), site in the top 10% percentile of predictability (1, top), and bottom 10% percentile (3, bottom). **E**. Same as **D** for monkey D. **F**. Predicted neuronal activity versus actual neuronal activity in response to checkerboard and moving bar presentations for monkey A sites 1, 2, and 3 shown in **D**. Each point represents activity in one 2×5 ms timepoint during the 400 ms presentation. **G**. Same as **F** for monkey D neuronal sites shown in **E. H**. Distribution of EV fraction in V1→V4 regressions of neural activity in response to visual stimuli in sites deemed visually reliable in monkey A (3 recording days, 30–44 V4 sites recorded per day; *n* = 132 total site recordings). **I**. Same as **H** for monkey D (2 recording days, 7–10 V4 sites recorded per day; *n* = 17 total site recordings). **J**. Distribution of EV faction for visually (purple) and non-visually (gray) reliable neurons in monkey V1 and V4. Both SNR over 2 and a split-half correlation value of over 0.8 were used to define a neuron to be visually reliable in monkeys L and A. In monkey D, a lower split-half correlation value of 0.6 was used to increase the site count. **K**. Distribution of EV fraction in V4→V1 regressions of neural activity in response to visual stimuli in sites deemed visually reliable in monkeys L (5 recording days, 537–592 V1 sites recorded per day; *n* = 2789 total site recordings), A (3 recording days, 251–381 V1 sites recorded per day; *n* = 989 total site recordings), and D (2 recording days, 8–10 V1 sites recorded per day; *n* = 18 total site recordings). **L**. Differences in EV fraction using different filtering methods to determine visually reliable neurons in macaque V4 and V1 across three monkeys (L, A, and D) V1 and V4 sites and two subsampled permutations of monkey L (to compare to site counts in monkey A and D). Split-half correlation value *thr* represents either 0.8 (monkeys L, A, and *L*_*A*_) or 0.6 (monkey D and *L*_*D*_. **M**. Head-to-head comparison of ridge regression EV and Poisson GLM EV on monkey MUAe for V1→V4 (left) and V4→V1 (right). Models share identical train/test folds, 25 ms bins, and temporal gaps; the Poisson GLM enforces non-negativity via a log link on raw MUAe (no baseline subtraction).

**Figure 2—figure supplement 2.**
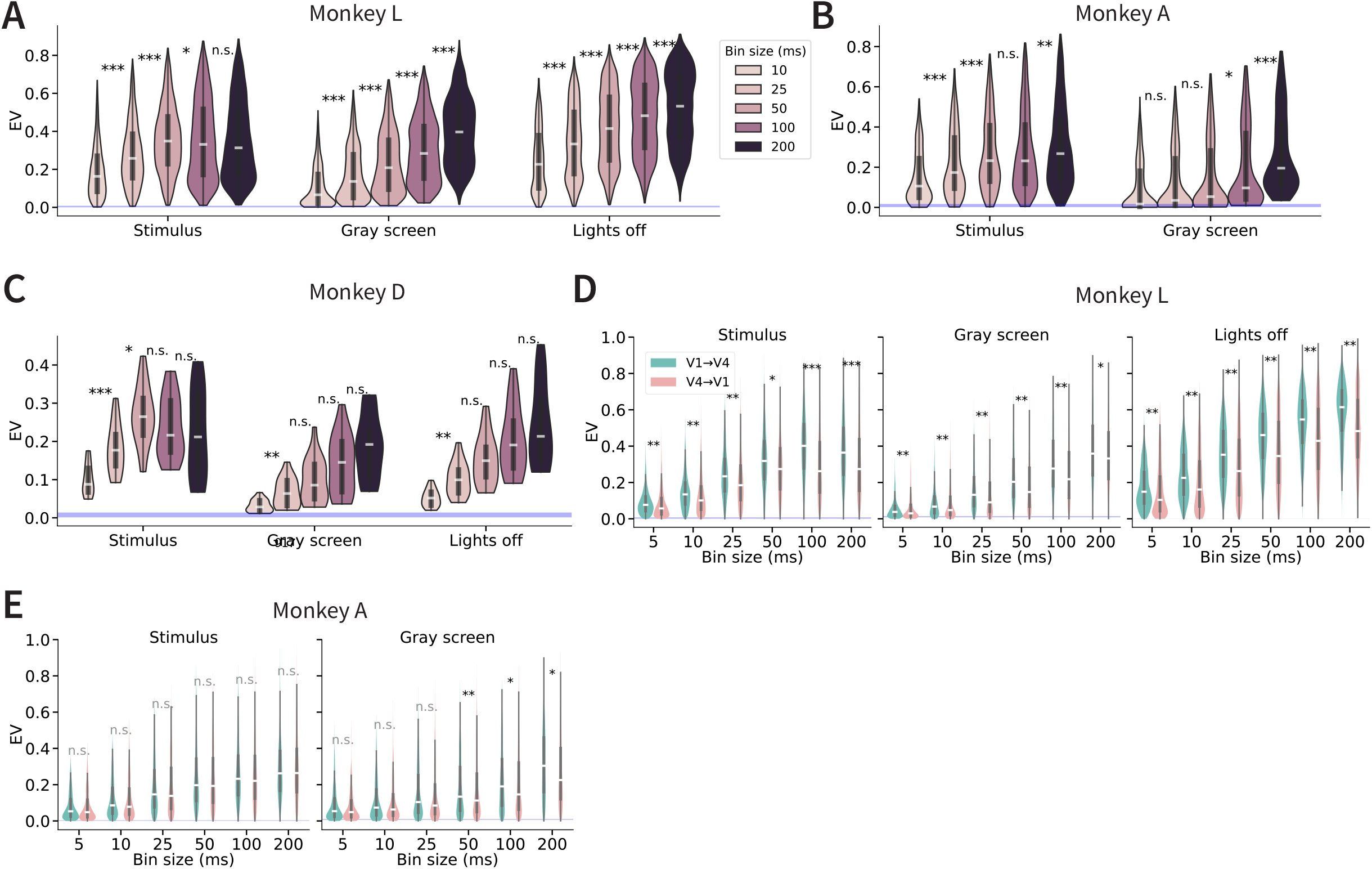
Inter-areal predictability across bin sizes. **A**. Distribution of EV fraction for V1↔V4 predictions in monkey L across stimulus, gray screen, and lights off conditions. Each violin plot corresponds to a different bin size (10–200 ms; color legend). **B**. same as **A** but for monkey A stimulus and gray screen conditions. **C**. Same as **A** but for monkey D. **D**. Same conditions as in **A**, but with sites randomly sub-sampled as in Fig. 3 to match both the number of sites and the distribution of split-half reliability values across the two prediction directions (V1→V4, teal; V4→V1, coral). **E**. Same as **D** but for monkey A stimulus and gray screen conditions.

**Figure 3—figure supplement 1.**
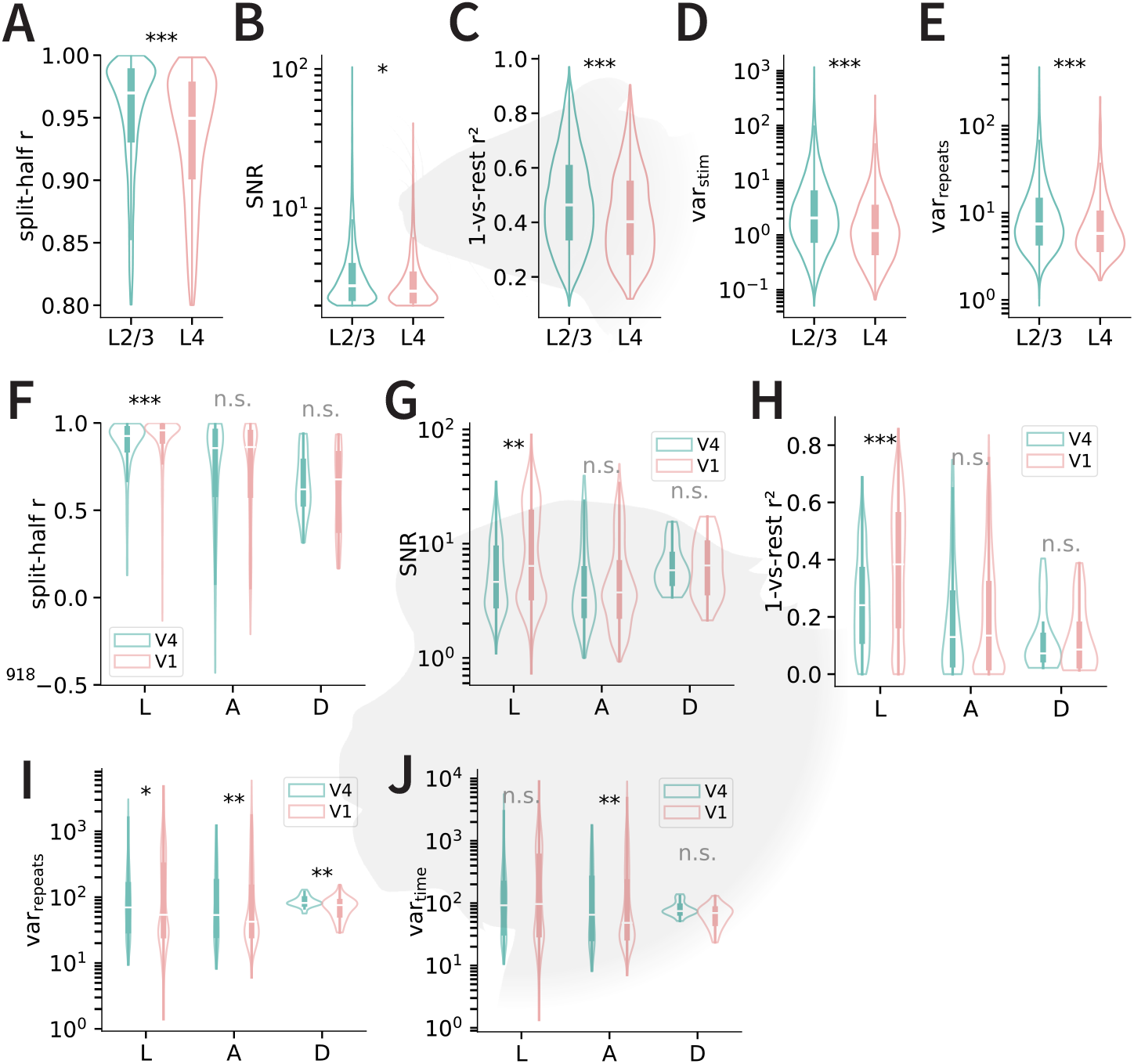
Neuronal property differences between areas in mouse and monkeys. **A–E**. Differences in self-consistency, SNR, 1-vs-rest squared, variance across stimuli, and variance across repeats between entire visually reliable neuronal populations in mouse L2/3 and L4. **F–J** Differences in self-consistency, SNR, 1-vs-rest squared, variance across time (within repeat), and variance across repeats (within timepoint) between entire visually reliable neuronal populations in monkey V1 and V4.

**Figure 3—figure supplement 2.**
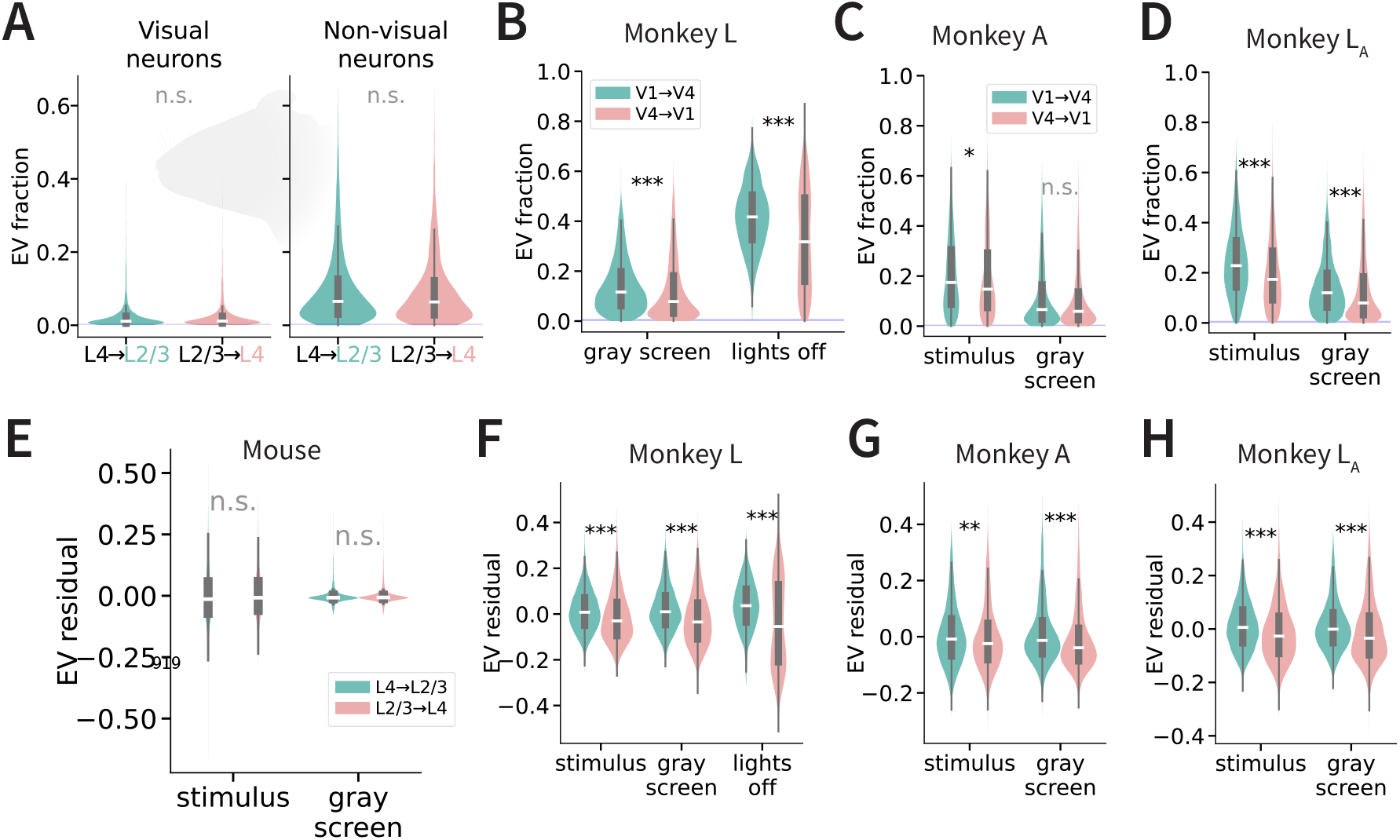
Differences in monkey inter-cortical predictability directions and lack of difference in mouse inter-laminar predictability directions are also seen in the absence of a stimulus. **A**. Differences in inter-laminar predictability directions in mouse neuronal activity during gray screen presentation in visual (left) and non-visual (right) neurons. **B**. Differences in inter-cortical predictability directions in monkey L during gray screen presentation and during lights off conditions. **C**. Differences in inter-cortical predicatibility directions in monkey A during stimulus and gray screen presentation. **D**. Same as **C** but for monkey L, after subsampling to match the number of sites in monkey A (*L*_*A*_). **E**. Differences in EV residuals after removing target-population covariates in mouse neural activity during stimulus and gray screen presentation (self-consistency, SNR, one-vs-rest *r*^2^, and variance metrics; for details see Methods). **F** Same as **E** but for monkey L EV residuals during stimulus, gray screen, and lights off conditions. **G**.Same as **E** but for monkey A EV residuals during stimulus and gray screen conditions. **H**. Same as **G** but for monkey L, after subsampling to match the site counts of monkey A (*L*_*A*_).

**Figure 4—figure supplement 1.**
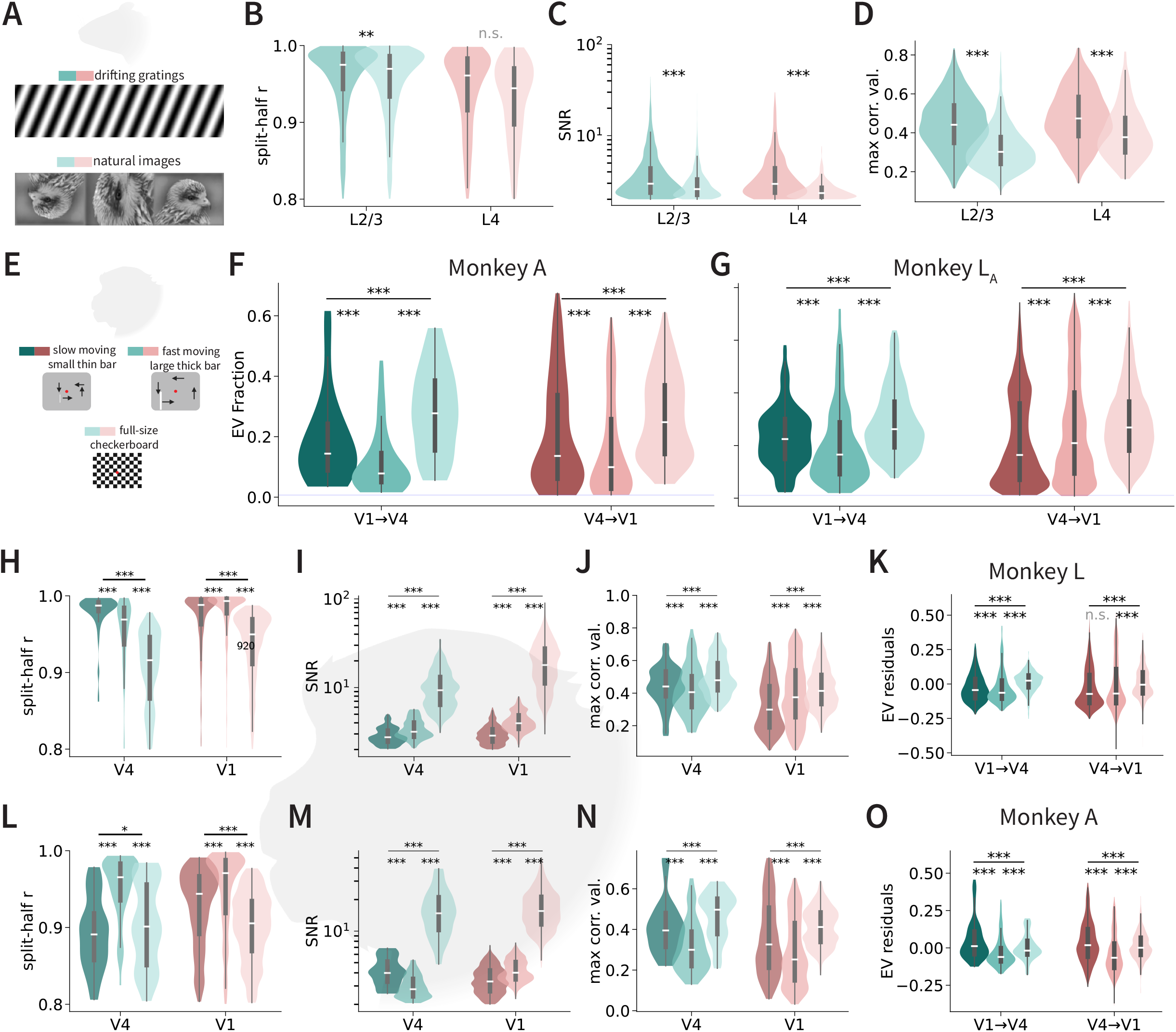
Neuronal activity properties for different stimulus types. **A**. Sample stimuli for mouse drifting grating and static natural images. **B-D**. Split-half correlation (**B**), SNR (**C**), and maximum correlation values (**D**) for each mouse layer and stimulus type (see color scheme in **A**). **E**. Sample stimuli for monkeys: full-size checkerboard, slow and fast moving bars. **F**. Across-area predictability for each stimulus type (dark: slow bars, medium: fast bars, light: checkerboard) and direction for monkey A. **G**. Same as **F** but for subsampled monkey L (*L*_*A*_). **H-I**. Same as **B-D** for monkey L V1 and V4 (see color map for each stimulus condition in **E**). **K**. EV *residuals* for monkey L after regressing, within direction, on SNR, split-half reliability, variance across time (within stimulus), and variance across trials (within timepoint). **L–N**. Same as **B-D** but for monkey A V1 and V4. **O**. Same as **K** but for monkey A.

**Figure 5—figure supplement 1.**
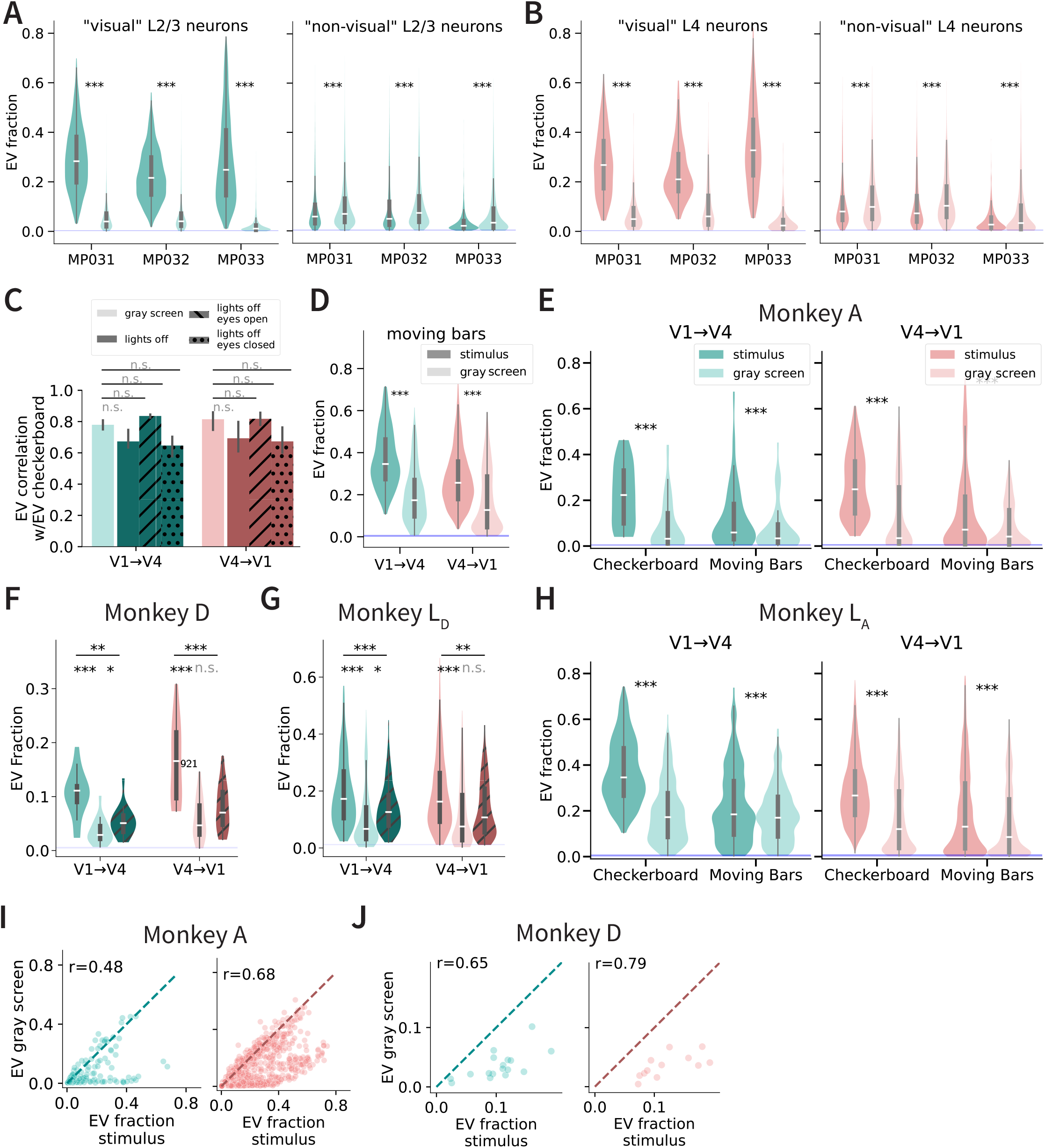
Comparing stimulus presentation vs. gray screen activity predictions in mouse and monkey. **A**. Differences in inter-laminar predictability between stimulus presentation and gray screen presentation neuronal activity in L4→L2/3 predictions across the three different mice (MP027 did not undergo gray screen presentation recordings). Left: visually reliable L2/3 neurons. Right: non-visually reliable L2/3 neurons. **B**. Same as **A**, but for mouse L2/3→L4 predictions. **C**. Correlation coefficient values between checkerboard presentation and gray screen and lights off conditions in monkey inter-cortical predictability. **D**. Differences in inter-cortical predictability between moving bar presentation and gray screen activity in monkey L V1→V4 and V4→V1 predictions (paired permutation test). **E**. Differences in inter-cortical predictability between stimulus and gray screen presentation across both checkerboard and moving bar stimuli in monkey A. **F**. EV differences in inter-cortical predictability between stimulus, gray screen, and lights off conditions in monkey D. **G**. Same as **F** but for monkey L subsampling sites to match the number of sites in monkey D (*L*_*D*_). **H**. Same as **E** but for monkey A, after subsampling sites to match the number of sites in monkey A (*L*_*A*_) **I**. Correlation between EV in responses to gray screen (y-axis) versus stimulus presentation (x-axis) in monkey A visually reliable neurons (V1→V4:left, green; V4→V1:right, coral). The diagonal line represents the line of equality (y=x). *r* is the Pearson’s correlation coefficient. **J**. Same as **I** for monkey D.

**Figure 5—figure supplement 2.**
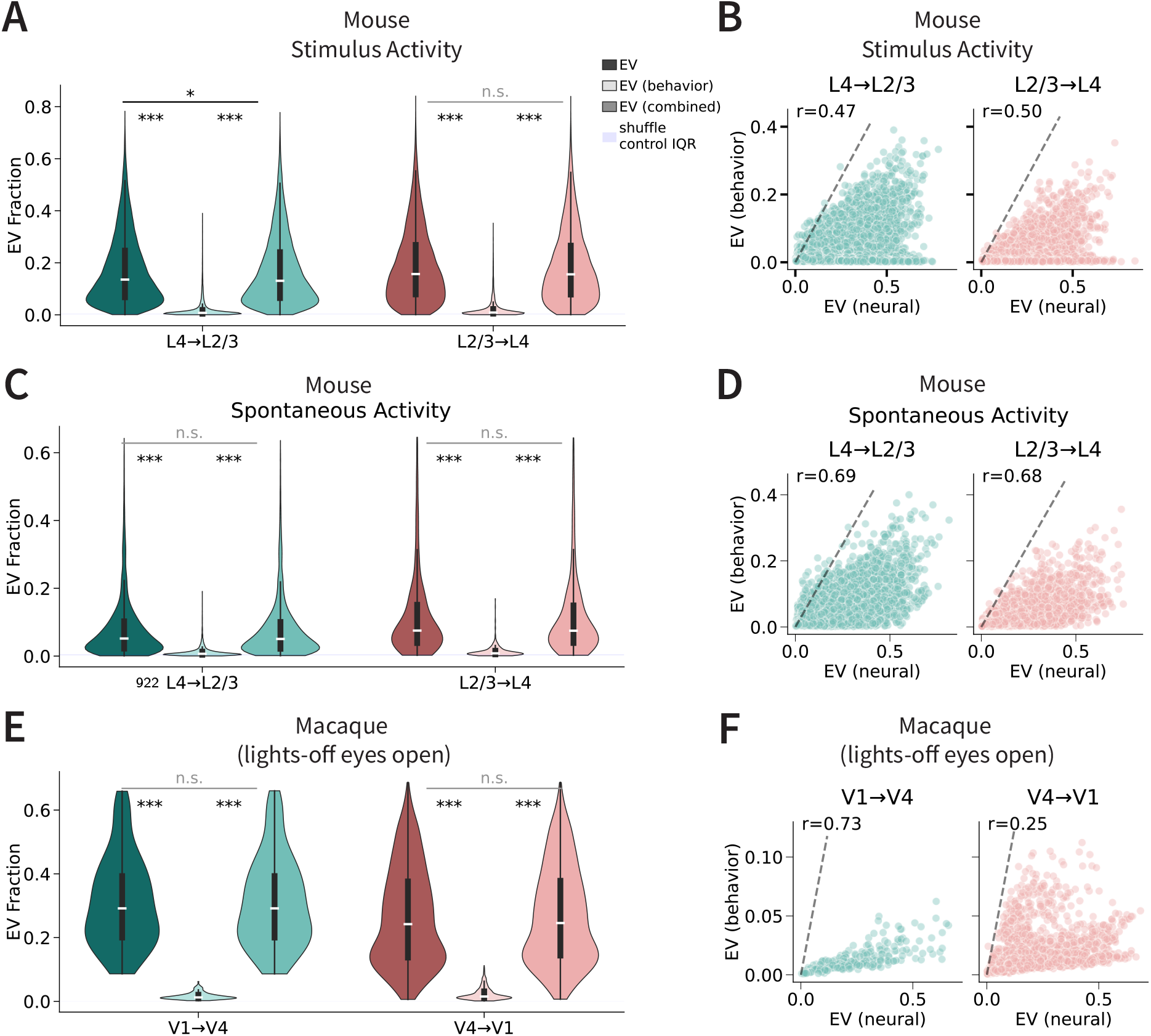
Behavioral contributions to inter-areal predictability in mouse and monkey. **A**. Distribution of EV fraction for neural-only (dark colors), behavior-only (face-motion SVD and running speed; light colors), and combined models (medium colors) in mouse L4→L2/3 and L2/3→L4 predictions during stimulus activity. **B**. Scatter plots comparing EV from behavior-only models (y-axis) versus neural-only models (x-axis) for mouse L2/3 response prediction (left) and mouse L4 response prediction (right). **C–D**. Same as **A–B**, but for spontaneous activity in mouse. **E**. Distribution of EV fraction for neural-only, pupil-only, and combined models in monkey L V1→V4 and V4→V1 predictions during resting state with eyes open. **F**. Scatter plots comparing EV from behavior-only models (y-axis) versus neural-only models (x-axis) for monkey V4 response prediction (left) and monkey V1 response prediction (right). The dashed line represents the line of equality (y=x).

**Figure 6—figure supplement 1.**
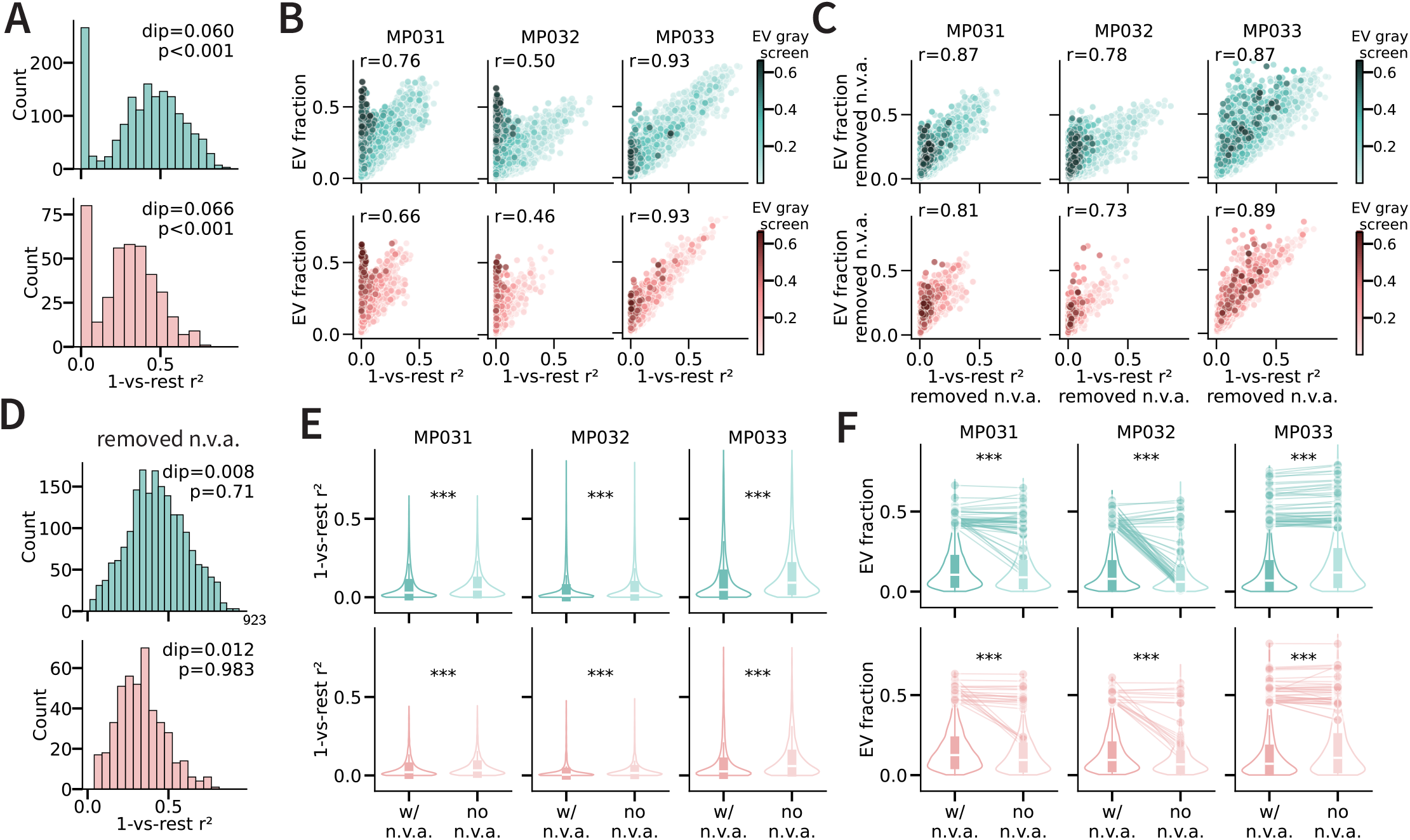
Properties of visual and nonvisually active neurons in mice. **A**. Bimodal distribution of 1-vs-rest squared correlation values in highly predictable L2/3 (top) and L4 (bottom) neurons (*EV* > 0.4). Hartigan’s dip test was applied to test bimodality (top right corner). **B**. Relationship between 1-vs-rest squared correlation and EV fraction in L2/3 (top, green) and L4 (bottom, coral) neurons in three mice (columns). **C**. Same as **B** after projecting out “non-visual ongoing” activity (***Stringer et al., 2019a***), see text for details. **D**. Distribution of 1-vs-rest squared correlation values in highly predictable L2/3 (top) and L4 (bottom) neurons (*EV* > 0.4) after projecting out “non-visual ongoing” activity. **E**. One-vs-rest squared correlation values when including (left) and not including (right) non-visual ongoing activity in each mouse (columns). * denote paired permutation test. **F**. EV fraction when including (left) and not including (right) non-visual ongoing activity dimensions in L2/3 (top, green) and L4(bottom, coral) in each mouse (columns). Sample subset of neurons with initial prediction values of over 0.4 visualized with lineplots. * denote paired permutation test.

**Figure 6—figure supplement 2.**
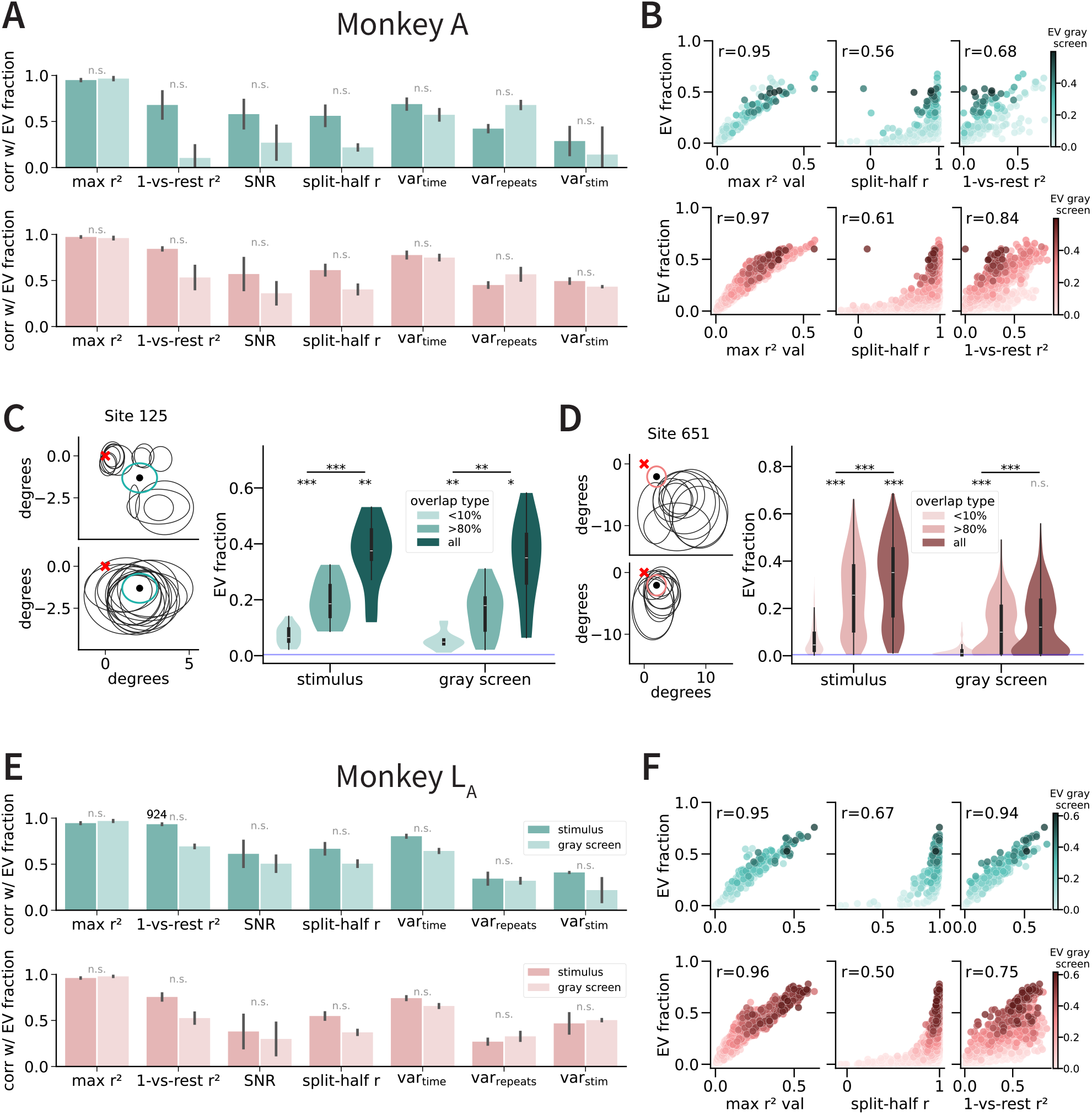
Influence of SNR, variance, and receptive field overlap is consistent across monkeys L and A. **A**. Correlation between different properties with the predictability of V4 (green) and V1 (coral) neuronal responses during the presence (dark color) or absence (light color) of visual stimulus in monkey A. **B**. Relationship between the EV fraction and three neuronal properties in sites for monkey A V4 (green) and V1 (coral) responses to visual stimuli. Hue represents the degree of predictability for the same sites during gray screen presentations (see color scale on right). **C**. Left, top: Receptive fields of one sample V4 neuronal site in monkey A (green circle, array 2 electrode 187) and 14 randomly selected V1 neuronal sites as predictors (black circles), constrained on sites that share less than 10% receptive field overlap with the V4 site. Bottom: Receptive fields of the same neuronal site 125 and 14 randomly selected V1 neuronal sites used as predictors, constrained on sites that share at least 80% receptive field overlap with the V4 site. Right: EV fraction of V4 neural activity (*n* = 18 site recordings per activity type) using 14 V1 predictor sites with less than 10% RF overlap (light green), 14 predictor sites with at least 80% RF overlap (green), and all predictors (dark green). Predictions were computed for recordings in response to the stimulus presentation (sliding bars and full-field checkerboard images) and gray screen presentation. **D**. Bottom and top left: Same as **F** but for monkey A sample V1 site 651. Right: Same as **C**, but for V1 (*n* = 378 site recordings per activity type). Instead of 14, 10 prediction sites were used to predict V1 due to low sample count of V4 that fulfilled both types of RF overlap percentages. **E–F**. Same as **A–B** but for monkey L after subsampling sites to match the number of sites in monkey A (*L*_*A*_).

**Figure 6—figure supplement 3.**
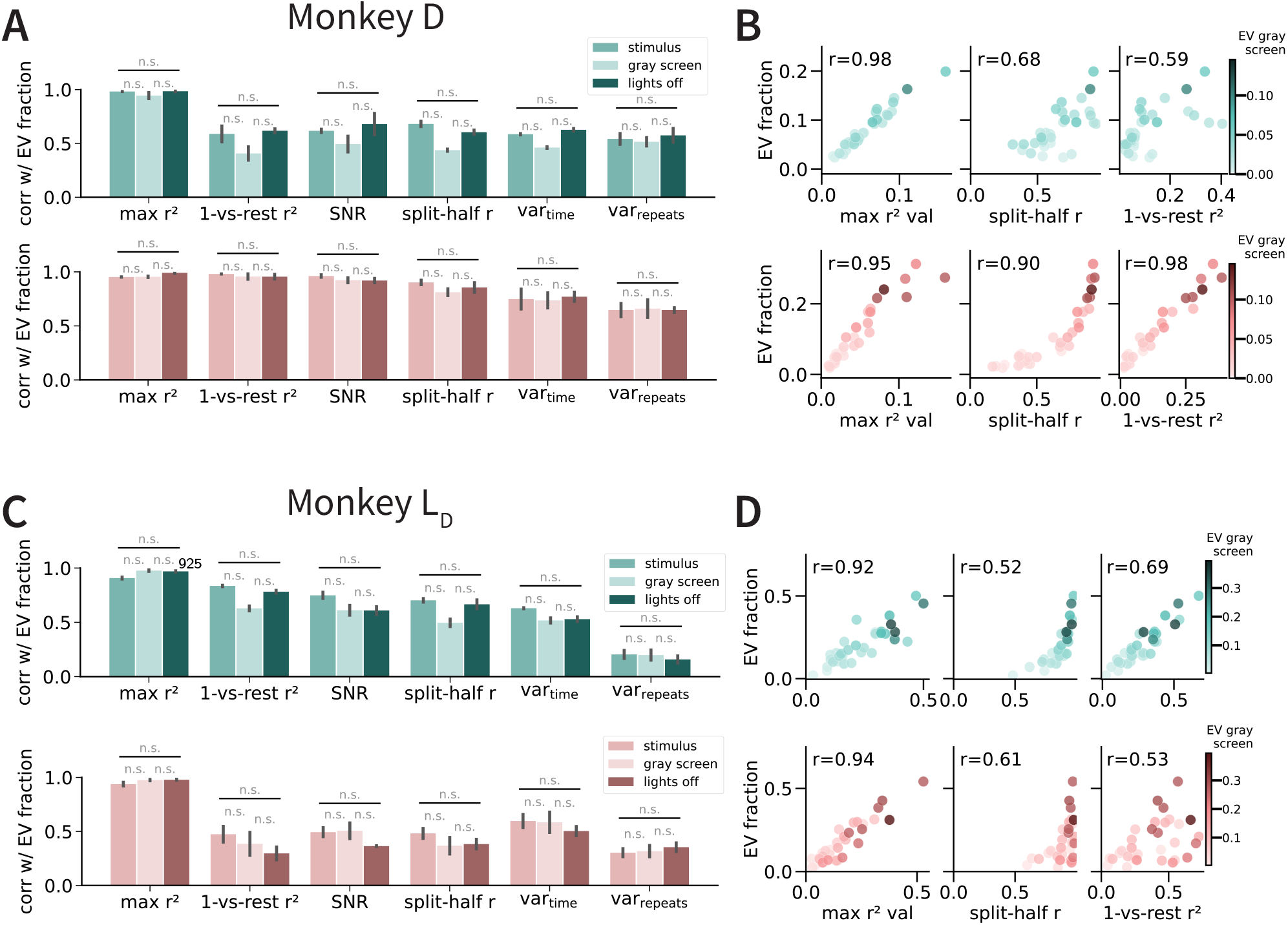
Influence of SNR, variance, and split-half reliability is consistent across monkeys L and D. **A**. Correlation between different properties with the predictability of V4 (green) and V1 (coral) neuronal responses during the presence (dark color) or absence (light color) of visual stimulus in monkey D. **B**. Relationship between three properties and their predictability in sites for monkey D V4 (green) and V1 (coral) responses from stimuli presentations. Hue represents the degree of predictability for the same sites during gray screen presentations. **C–D**. Same as A–B but for subsample monkey *L*_*D*_.

**Figure 7—figure supplement 1.**
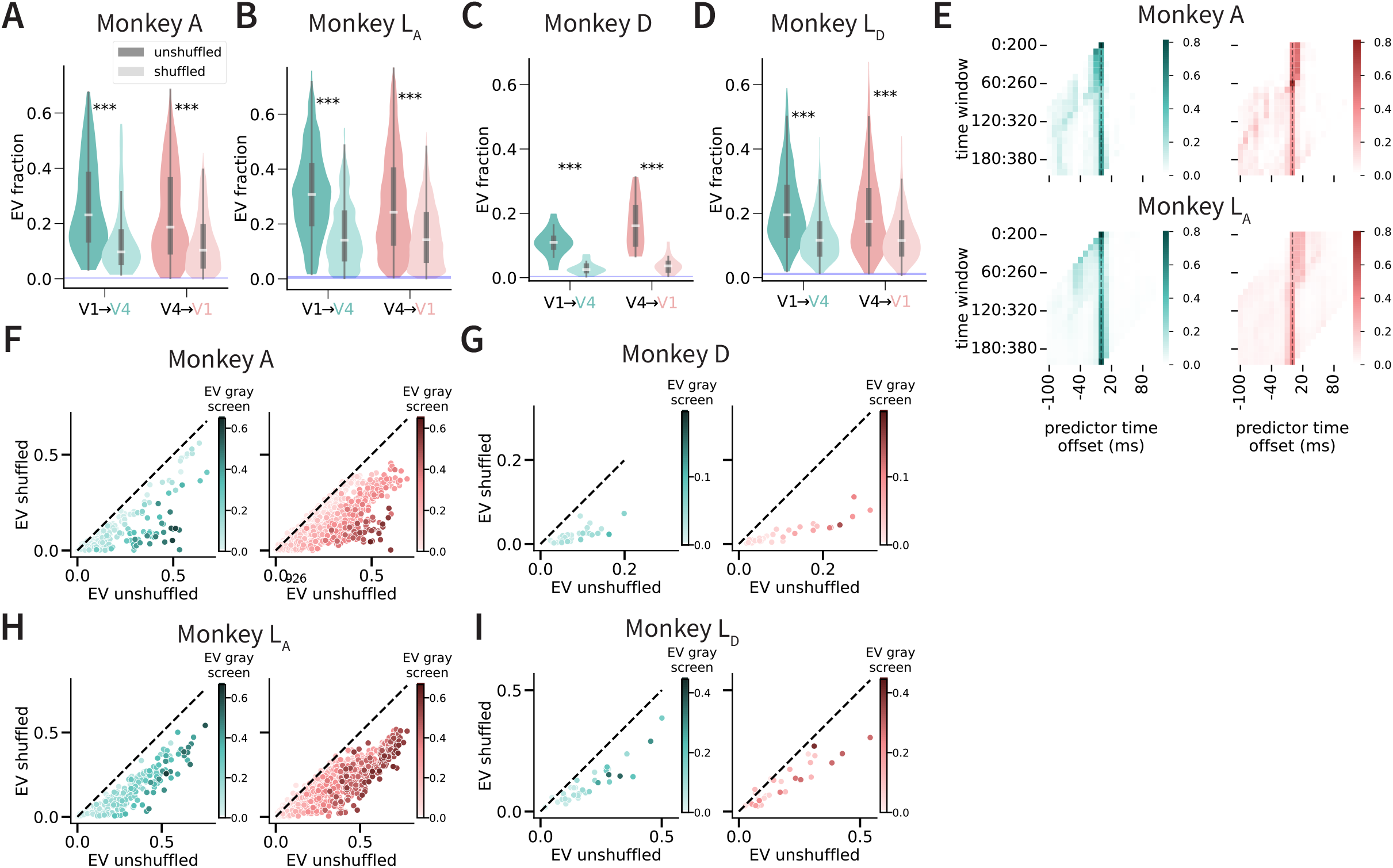
Time-dependent effects on EV predictability in monkeys A and D. **A–D** Distribution of EV fraction in unshuffled (dark) and shuffled (light) trial-repeat activity in monkey A, monkey L subsampled to match the number of sites in monkey A (*L*_*A*_), monkey D, and monkey L subsampled to match the number of sites in monkey D (*L*_*D*_). * denote paired permutation tests. **E**. Time offset prediction results across both V1→V4 (left, green) and V4→V1 (right, coral) prediction directions in monkeys A (top) and subsampled *L*_*A*_ (bottom). Each square corresponds to the fraction of neuronal sites whose neural activity was best predicted during that offset period and time window. **F–I**.Relationship between shuffled (y-axis) and unshuffled (x-axis) trial repeat EVs in V1 → V4 (left, green) and V4 → V1 (right, coral) directions in monkeys A, subsampled *L*_*A*_, D, and subsampled *L*_*D*_. Hue represents EV fraction during gray screen activity (see color scale on right).

**Figure 7—figure supplement 2.**
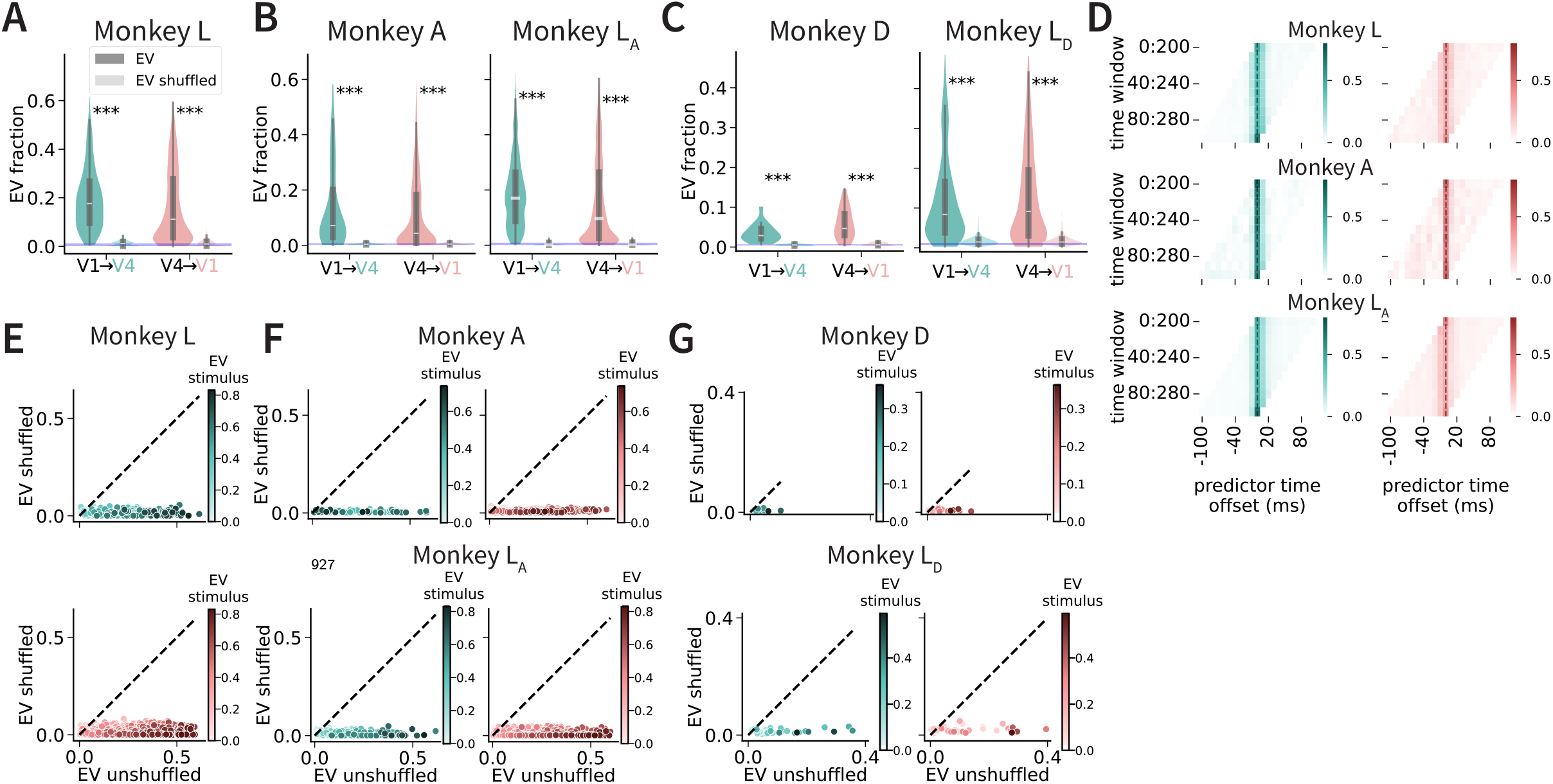
Time-dependent effects on EV predictability during spontaneous conditions. Distribution of EV fraction for unshuffled (dark) and shuffled (light) trial-repeat activity during gray screen presentations (paired permutation test) in monkeys L, A, subsampled *L*_*A*_, D, and subsampled *L*_*D*_. * denote paired permutation tests. **D**. Time offset prediction results across both V1→V4 (left, green) and V4→V1 (right, coral) prediction directions during gray screen presentations in monkeys A (top) and subsampled *L*_*A*_ (bottom). Each square corresponds to the fraction of neuronal sites whose neural activity was best predicted during that offset period and time window. **E–G**.Relationship between shuffled (y-axis) and unshuffled (x-axis) trial repeat EVs in V1 → V4 (left, green) and V4 → V1 (right, coral) directions in monkeys L (**E**), A & subsampled *L*_*A*_ (**F**), and D & subsampled *L*_*D*_ (**G**). Hue represents EV fraction during stimulus activity (see color scale on right).

**Figure 7—figure supplement 3.**
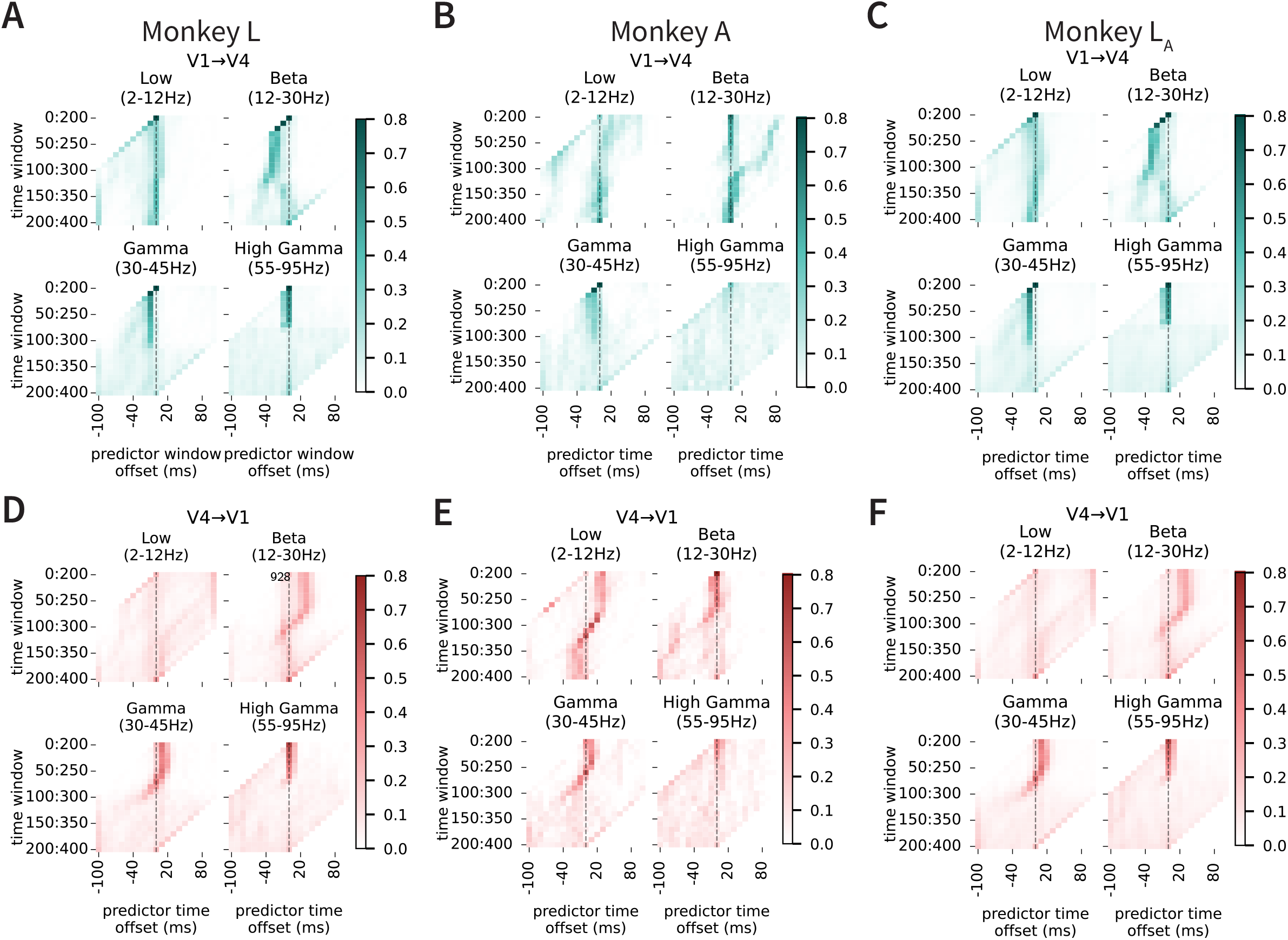
Time-dependent effects on EV predictability across LFP bands in monkey V1/V4. **A–C**. Time offset prediction results across both V1→V4 (left,green) and V4→V1 (right, coral) prediction directions in monkeys L, A, and *L*_*A*_. Columns correspond to band-limited LFP amplitude (Hilbert envelope) from Low (2–12 Hz), Beta (12–30 Hz), Gamma (30–45 Hz), and High-gamma (55–95 Hz). LFP preprocessing included removal of narrow line artifacts (notch filter at 50 Hz and harmonics), band-pass filtering, and Hilbert amplitude extraction; envelopes were z-scored per unit. The dashed vertical line marks zero offset. **D–F**. Same as **A–B**, but predicting V1 from V4.

